# Overcoming the eIF2α Brake in Human Cell-Derived Translation Systems

**DOI:** 10.1101/2025.11.16.688697

**Authors:** Nikolay A. Aleksashin, Rohan R. Shelke, Tianhao Yin, Jamie H. D. Cate

## Abstract

Cell-free translation from human cells is a powerful platform for studying mammalian gene expression and building synthetic biology tools, but productivity is often curtailed by inhibitory phosphorylation of eIF2α on residue Ser52. Here we systematically explored complementary strategies to bypass this initiation block across editable and hard-to-edit human cell types. In Expi293F suspension cells, precise genome editing of *EIF2S1* to block Ser52 phosphorylation (eIF2α-S52A) produced high-activity extracts. Genetic knockout of *EIF2AK2* (PKR)–the principal eIF2α kinase engaged in eIF2α phosphorylation in Expi293F lysates–also improved translation, further establishing eIF2α phosphorylation as the dominant bottleneck in Expi293F translation extracts. Because genome editing is impractical in many contexts including primary human cells, we also implemented expression-based rescue of eIF2α phosphorylation: stable piggyBac integration of truncated GADD34 (*PPP1R15A*) and K3L, a viral eIF2α decoy, under control of a Tet-inducible promotor in induced pluripotent stem cells (iPSCs) and primary human fibroblasts. After differentiating engineered KOLF2.1J iPSCs into cardiomyocytes, we found that stable GADD34/K3L expression increased translation output in cardiomyocyte translation extracts. Using the piggyBac expression system in primary fibroblasts also resulted in improved translational output. Together these data pinpoint eIF2α phosphorylation as the key barrier to robust translation in human cell translation extracts. They also show that editing eIF2α or removing PKR is optimal where genome editing is feasible, while providing a portable GADD34/K3L expression cassette enables production of translationally active lysates from systems where genome editing is challenging or not possible.

## Introduction

Cell-free translation systems provide a versatile framework for dissecting the mechanisms of gene expression and reconstituting complex regulatory events, and enable diverse applications in synthetic biology (1). *In vitro* translation (IVT) reactions recapitulate essential steps of mRNA decoding and nascent polypeptide synthesis under defined biochemical conditions, offering an experimentally accessible counterpart to cellular protein synthesis. Among these systems, those derived from mammalian cells are uniquely positioned to capture features of the human translational apparatus (2, 3), including its specialized ribosomal proteins, species-specific features of translation factors, and the post-translational modification landscape (e.g. phosphorylation of initiation/elongation factors) (4–6). Beyond fundamental research, mammalian IVT systems hold growing promise in biotechnology and personalized medicine (7, 8). Because they maintain key components responsible for species-specific post-translational modifications and native chaperone interactions, such extracts can additionally support synthesis of eukaryotic proteins that are challenging to express in bacterial or yeast systems (9). Moreover, cell type-specific human lysates can serve as *in vitro* proxies for modeling translation control, disease-relevant pathways, or drug responses in a biochemical format. The prospect of producing lysates from patient-derived or lineage-specific cells further fuels interest in precision biochemical assays of human translation regulation.

Despite this potential, a central limitation of human cell–derived IVT systems is their relatively low translational efficiency compared to optimized systems from *E. coli*, wheat germ, or rabbit reticulocyte lysates (7, 10, 11). Low translational yields reflect both the biochemical complexity of mammalian translation and unintended activation of regulatory checkpoints during extract preparation. Residual stress kinase activity during lysis can lead to phosphorylation of translation factors such as eIF2α, which suppresses global initiation (4, 5, 12). In eukaryotes, global protein synthesis is dynamically regulated through reversible phosphorylation of translation factors to tune initiation and elongation in response to cellular state (**Fig. 1A**) (13). Phosphorylation of eIF2α at Ser52 converts eIF2 from an active initiation factor to a dominant inhibitor of its guanine nucleotide exchange factor eIF2B, thereby depleting the available ternary complex and suppressing translation (13–15) (**Fig. 1B**). This pathway is central to the integrated stress response (ISR), where diverse upstream kinases (PKR, GCN2, PERK, HRI) sense stresses such as dsRNA, ER stress, amino acid insufficiency, or heme limitation, and relay the signal via eIF2α phosphorylation (16–18). In parallel, elongation efficiency is modulated through phosphorylation of eukaryotic elongation factor 2 (eEF2) at Thr57 by eEF2 kinase (eEF2K) (19–22), which inhibits ribosomal translocation along the mRNA and transiently slows peptide elongation (**Fig. 1C**). This modification provides a rapid, reversible means of reducing translational throughput during cellular stress, energy limitation, or nutrient deprivation (21, 23, 24). Like eIF2α phosphorylation, eEF2 phosphorylation is widely used to tune protein synthesis rates across physiological and stress conditions, though its relative contribution to translation control in cell-free lysates remains poorly understood (25).

**Figure 1.**
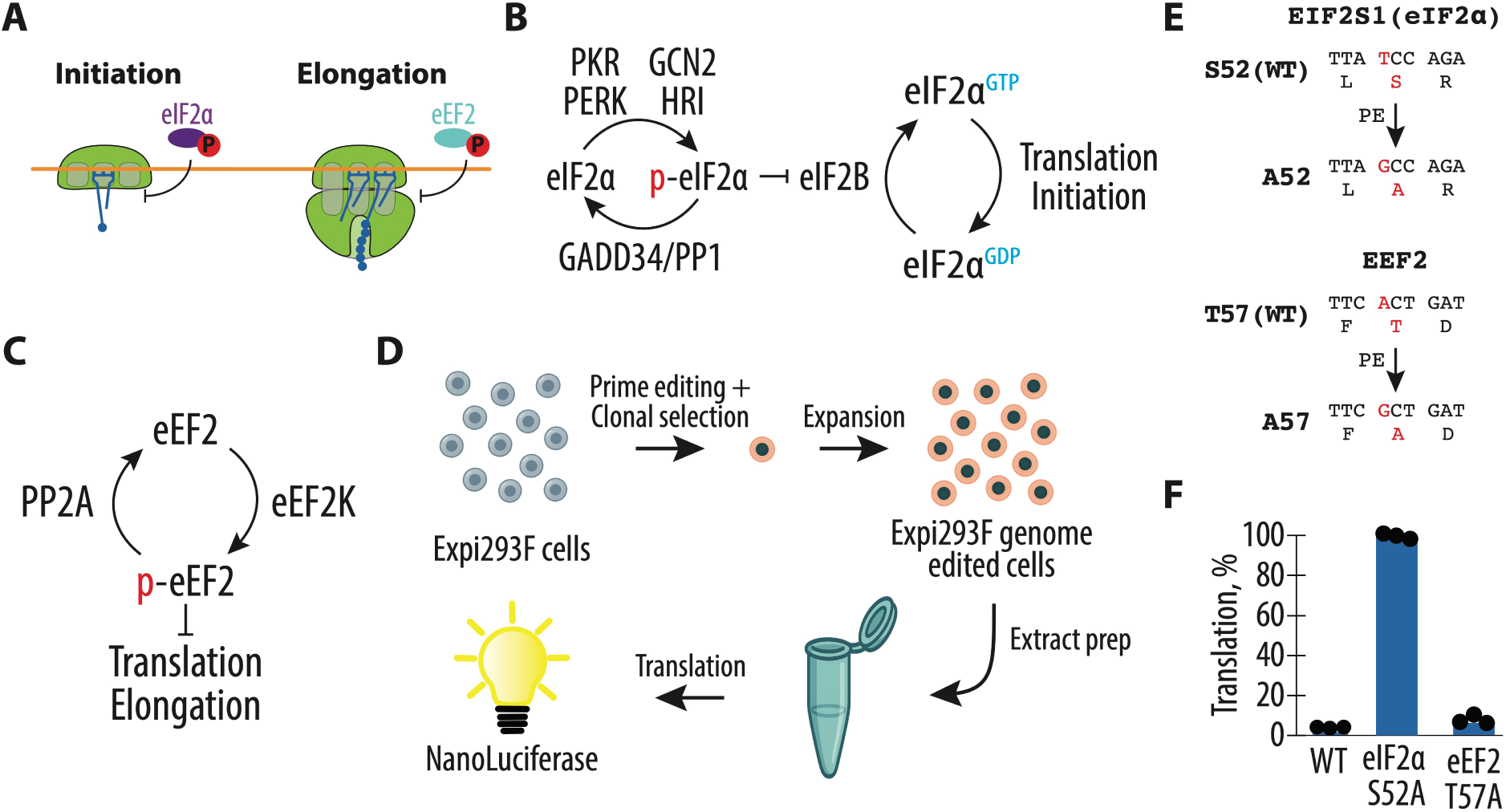
Blocking phosphorylation of eIF2α, but not eEF2, enhances translational efficiency in human Expi293F extracts. **(A)** Left: Phosphorylation of eIF2α on Ser52 converts eIF2 into an inhibitor of its guanine nucleotide exchange factor (eIF2B), blocking ternary complex regeneration and suppressing translation initiation. Right: Phosphorylation of eEF2 on Thr57 by eEF2 kinase (eEF2K) prevents ribosomal translocation, thereby slowing peptide elongation. **(B)** Phosphorylation of eIF2α (Ser52) by stress-activated kinases (PKR, GCN2, PERK, HRI) converts active eIF2 into an inhibitor of its guanine nucleotide exchange factor eIF2B, thereby preventing ternary complex formation and suppressing global initiation. **(C)** The eEF2 cycle: eEF2K phosphorylates eEF2 at Thr57, which prevents ribosomal translocation; dephosphorylation by PP2A restores elongation. **(D)** Illustration of the experimental workflow for lysate generation and *in vitro* translation (IVT). Suspension Expi293F cells were harvested, lysed under native conditions, and the resulting extracts programmed with a Nanoluciferase (NanoLuc) reporter mRNA to quantify translational efficiency. **(E)** Schematic representation of human Expi293F cells engineered by prime editing (PE) to introduce phospho-null substitutions in *EIF2S1* (eIF2α S52A) and *EEF2* (eEF2 T57A). All amino acid residue numbers correspond to the human reference sequences according to the UniProt database, entries P05198 and P13639 for *EIF2S1* and *EEF2,* respectively. **(F)** Comparison of translational output in IVT reactions programmed with NanoLuc mRNA using extracts prepared from wild-type Expi293F cells or genome-edited Expi293F eIF2α-S52A and eEF2-T57A lines. All experiments were performed in biological triplicates.

In mammalian cell lysates, even minor unintended activation of eIF2α kinases during extract preparation can carry over into the IVT reaction, causing spontaneous accumulation of phospho-eIF2α and drastic suppression of translation output (4, 5, 10, 12, 26–28). To bypass or reverse eIF2α phosphorylation in a controlled way, several strategies have been explored. One route is to supply the *in vitro* translation reaction with the regulatory subunit GADD34, which targets phosphatase PP1 to dephosphorylate eIF2α and serves as a built-in feedback loop in cellular stress regulation (10, 27–30). Another is the use of viral decoy proteins, such as vaccinia K3L, which mimic eIF2α and act as competitive substrates to sequester kinases like PKR (31–34). GADD34 and K3L addition or co-expression was used in prior work to suppress eIF2α phosphorylation and produce extracts with higher translational yields (5, 10, 27, 29, 35, 36) (**Fig. 1B**). Although these strategies have shown promise in HEK293T or HeLa-based systems, a systematic comparison across mammalian cell types and extract preparations is lacking. It is unclear which kinase(s) dominate eIF2α phosphorylation in different lysates, how effective editing or knockout strategies may be in relieving suppression, and how inducible expression of dephosphorylation factors might scale to diverse cell types including iPSC-derived cell types and primary human cells.

Here, we sought to systematically define molecular constraints on translational output and to implement complementary genetic and expression-based strategies to overcome them. We focus on inhibitory phosphorylation of eIF2α as a primary bottleneck. We show that editing *EIF2S1* to prevent eIF2α phosphorylation in Expi293F cells yields extracts with significantly enhanced activity, that partial deletion of PKR also relieves suppression, and that inducible co-expression of GADD34 and K3L can improve translation even in systems refractory to genome editing, such as iPSC-derived cardiomyocytes and primary fibroblasts. Together, our results provide a versatile and robust foundation for generating high-efficiency, human cell–derived translation systems across cell types.

## Materials and Methods

### Cloning of pSBtet-RP-PEMAX

The Sleeping Beauty transposon vector pSBtet-RP-PEMAX was constructed using the Addgene plasmid pSBtet-GP (37) (Addgene #60497; gift from the Ingolia lab) as the backbone. The vector was linearized with the SfiI restriction enzyme (New England Biolabs) according to the manufacturer’s instructions. The PEmax–P2A–dnMLH1 insert was amplified by PCR from the template plasmid pCMV-PEmax-P2A-hMLH1dn (38) (Addgene #174828) using Q5 High-Fidelity DNA Polymerase (New England Biolabs) and the following primers:

Forward: 5′CCACTTCCTACCCTCGAAAGGCCTCTGAGGCCACCATGAAACGGACAGCCGACGG AAGCG-3′

Reverse: 5′TCATGTCTATCGATGGAAGCTTGGCCTGACAGGCCTTAGAAAACCTTATAAAGGTC GGGC-3′

The PCR product was verified by agarose gel electrophoresis, digested with DpnI overnight at 37 °C to remove template DNA, and purified using the DNA Clean & Concentrator-5 Kit (Zymo Research). Gibson Assembly was performed using the NEBuilder HiFi DNA Assembly Master Mix (New England Biolabs) following the manufacturer’s protocol. The resulting construct was sequence-verified by full plasmid sequencing and has been deposited at Addgene (Plasmid #249032).

### Expi293F Cell Culture

Expi293F cells (from UC Berkeley cell culture facility) were maintained in Expi293F Expression Medium and cultured according to the manufacturer’s instructions. Cells were propagated in vented Erlenmeyer flasks at 37 °C in a humidified incubator with 8% CO₂ and continuous shaking at 125 rpm. For experiments requiring adherent growth (e.g., genome editing and transfection assays), Expi293F cells were adapted to adherent conditions by passaging in DMEM medium supplemented with 10% Tet-system approved FBS (Gibco) and plated onto standard tissue culture–treated plasticware. Cells were passaged every 3–4 days to maintain exponential growth.

### Generation of Expi293F cells with a stably integrated prime editing construct

A stable Expi293F cell line expressing a prime editing construct (pSBtet-PuroR/dTomato-PEMax) was generated using the Sleeping Beauty transposon system (37). Cells were nucleofected with 100 ng of pSB100X transposase plasmid and 1000 ng of pSBtet-PuroR/dTomato-PEMax plasimid using the Amaxa 4D-Nucleofector X Unit (Lonza) and SF cell line kit according to the manufacturer’s instructions. After two days of recovery in antibiotic-free medium, cells were subjected to puromycin selection (1 µg/mL; Gibco) for 5–7 days in DMEM medium (Gibco) supplemented with 10% Tet-system approved FBS (Gibco). The efficiency of selection was monitored visually by observing dTomato fluorescence and cell viability. Following selection, stable populations were expanded and maintained in Expi293F Expression Medium under standard culture conditions.

### Construction of epegRNA and sgRNA Expression Plasmids

The epegRNA and sgRNA expression vectors were amplified by PCR from pU6-tevopreq1-GG-acceptor (Addgene, 174038) and BPK1520 plasmid (a gift from the Dirk Hockemeyer lab) respectively using Q5 High-Fidelity DNA Polymerase (New England Biolabs) following the manufacturer’s instructions. All oligonucleotides used for cloning of epegRNA and sgRNA constructs are listed in **Table S1**. All epegRNA and sgRNA sequences for prime editing experiments were designed using Primedesign (39). The reactions were cycled with an initial denaturation at 98 °C for 30 s, followed by 30 cycles of 98 °C for 10 s, 67 °C for 30 s, and 72 °C for 66 s, and a final extension at 72 °C for 2 min. PCR products were analyzed by agarose gel electrophoresis (1% agarose in 1× TAE buffer, 110 V for 20 min) to confirm the expected product size (2.2 kb). Verified PCR products were digested with DpnI (New England Biolabs) in 1× CutSmart Buffer overnight at 37 °C to remove template plasmid DNA, followed by purification using the DNA Clean & Concentrator-5 Kit (Zymo Research).

Gibson assemblies were performed using NEBuilder HiFi DNA Assembly Master Mix (New England Biolabs) to insert target-specific variable oligonucleotides into the linearized epegRNA or sgRNA vectors. For epegRNA plasmids, reactions (10 μL each) contained 1× HiFi Master Mix, 30 ng of vector PCR, and 0.5 pmol each of Constant Oligo 1 and Constant Oligo 2, with 0.5 pmol of the desired variable oligo (all oligos were diluted in 1× NEBuffer 2.1 buffer). Reactions were incubated at 50 °C for 1 h. For sgRNA plasmids, Gibson reactions (10 μL total) included 1× HiFi Master Mix, 1 μL of sgRNA vector (30 ng/μL), and 1 μL of each variable oligo (1 μM stock concentration in 1x NEBuffer 2.1 buffer). Reactions were incubated at 50 °C for 1 h. Following assembly, 5 μL of the reaction was transformed into chemically competent *E. coli* DH5alpha (NEB) by heat shock at 42 °C for 40 s, recovered in 1000 μL SOC medium at 37 °C for 1 h with shaking, and plated on LB-ampicillin agar plates. Individual colonies were inoculated into 2 mL LB medium with ampicillin and grown overnight at 37 °C. Plasmid DNA was purified using the Zyppy Plasmid Miniprep Kit (Zymo Research, D4036) and verified by sequencing.

### Prime Editing Experiments in Expi293F Cells

Prime editing was performed in Expi293F cells harboring the stably integrated pSBtet-PuroR/dTomato-PEMax construct described above. The workflow is schematically summarized in **Figure S1**. Briefly, following induction of PEmax expression with doxycycline (1 µg/mL; Takara) for 24 h, cells were nucleofected with the corresponding epegRNA and nicking sgRNA plasmids using the Amaxa 4D-Nucleofector X Unit (Lonza) and the SF Cell Line Nucleofection Kit, according to the manufacturer’s protocol. For each nucleofection, 2 × 10^4^ cells were mixed with 66 ng of epegRNA and 22 ng of sgRNA plasmid DNA in a final volume of 20 µL nucleofection buffer. After electroporation, cells were immediately transferred into pre-warmed DMEM medium (Gibco) supplemented with 10% Tet-system FBS (Gibco) and allowed to recover for 48 h.

After 72 h, cells were washed trice with PBS, pH 7.4 (Gibco) and incubated at 37 °C for 1 h in cell lysis buffer (10 mM Tris-HCl, pH 8.0 (Invitrogen), 0.05% SDS (Promega), 1 u/mL Proteinase K (New England Biolabs), followed by incubation at 80 C for 20 min to inactivate proteinase K. Regions of interest were amplifying by PCR with PrimeSTAR GXL DNA polymerase (Takara) according to the manufacture instructions. PCRs were done using the primers:

eIF2 Fwd: 5′-GCAAAATAAAAATTAAAGCTTGGTT-3′;

eIF2 Rev: 5′-GTCAATTATCAAGATCCTGATAAAC-3′;

eEF2 Fwd: 5′-GGTGGTTCCTTGGAGATT-3′;

eEF2 Rev: 5′-TCCCAGGTGTGACAGCCA-3′, respectively. Amplicons were purified using Zymo DNA concentrator kit and subjected to Plasmidsaurus amplicon sequencing. For clonal validation, edited cell populations were serially diluted into 96-well plates, expanded, and screened for the desired allelic modifications by sequencing. Verified clones were subsequently expanded, cryopreserved, and used for downstream biochemical assays.

### Generation of PKR and eEF2K Knockout Expi293F Cells

Gene knockouts of PKR and eEF2K were generated in Expi293F cells adapted to adherent culture conditions in DMEM media (Gibco) supplemented with 10% FBS (Gibco). Genome editing was performed using CRISPR/Cas9 ribonucleoprotein (RNP) complexes assembled from recombinant Cas9 nuclease (Thermo Fisher Scientific) and synthetic TrueGuide single guide RNAs (sgRNAs; Thermo Fisher Scientific) following the manufacturer’s instructions. The sgRNA target sequences were as follows: PKR: 5′-TTATGAACAGTGTGCATCGG-3′; eEF2K: 5′-GTGTCGAGTAGCACGTTCGG-3′. All sgRNAs and oligonucleotides for sequencing were designed using the Invitrogen TrueDesign Genome Editor.

Cells were seeded into six-well tissue culture plates 24 h prior to transfection to achieve ∼70–80% confluency at the time of transfection. RNP complexes were prepared immediately before use by combining Cas9 protein and sgRNA at a 1:1.2 molar ratio and incubating for 10 min at room temperature to allow complex formation. Transfection was performed using Lipofectamine CRISPRMAX reagent (Thermo Fisher Scientific) according to the manufacturer’s protocol, with each well receiving preformed RNP–lipid complexes diluted in Opti-MEM Reduced Serum Medium. After 48 h, editing efficiency was assessed in pooled cells using the T7 Endonuclease I assay (GeneArt Genomic Cleavage Detection Kit, Thermo Fisher Scientific).

Clonal populations were established by limiting dilution in filtered 50% conditioned/50% fresh DMEM medium and expanded individually over 2 weeks. Successful knockout clones were verified by Sanger sequencing of the targeted genomic region and by Western blot analysis of PKR and eEF2K protein levels. Saenger sequencing analysis were conducted using TIDE online software (40). PCRs for sequencing were done using corresponding primers: PKR Fwd: 5′-AGTCACACTTTTGCTCAAGGT-3′; PKR Rev: 5′-GGTTCCATGGCTTCAGCAAT-3′; eEF2K Fwd: 5′-GTAATGCACCGGGCCTTAAG-3′; eEF2K Rev: 5′-TGTGCCTGTGAGTGCAGATA-3′. Confirmed knockout lines were expand, adopted to the suspension culture conditions and maintained under standard Expi293F culture conditions (Gibco) at 37 °C in a humidified atmosphere with 8% CO₂.

### Preparation of Expi293F Cell Extracts

Cells were collected by centrifugation at 2000 × *g* for 15 min at 4 °C, and the supernatant was discarded. The pellet was washed once with ice-cold Dulbecco’s phosphate-buffered saline (DPBS) (Gibco), followed by centrifugation at 2000 × *g* for 15 min at 4 °C. Approximately 10⁸ cells typically yielded 0.2–0.3 mL of packed cell volume. The supernatant was removed, and the pellet was resuspended in ice-cold DPBS (∼2 mL per pellet). The suspension was divided equally into two 1.5 mL microcentrifuge tubes (∼1 mL per tube) and centrifuged again at 2000 × *g* for 5 min at 4 °C. The supernatant was discarded, and the pellet was resuspended in five cell volumes (∼0.5 mL per tube) of ice-cold Buffer A (10 mM HEPES–KOH, pH 7.5 (Thermo Scientific), 10 mM KCl (Ambion), 0.5 mM MgCl_2_ (Thermo Scientific) and incubated on ice for 20 min. The cells were then pelleted by centrifugation at 2000 × *g* for 5 min at 4 °C, and the supernatant was removed.

The resulting pellet was resuspended in one cell volume of ice-cold Buffer A (∼0.5 mL per 10^8^ cells) and lysed by passing the suspension through a 1 mL syringe fitted with a 26 G needle (BD PrecisionGlide, 26G × ½ [0.45 mm × 13 mm], Cat. No. 305111) approximately 10–15 times. Smooth, even strokes were applied to ensure uniform lysis while minimizing foaming and mechanical stress. Excessive handling was avoided to preserve extract activity. Lysates from parallel preparations were combined and centrifuged at 20, 000 × *g* for 10 min at 4 °C to remove cell debris and nuclei. The clarified supernatant (A₂₆₀ ≈ 50–60) was aliquoted (55 µL per tube), flash-frozen in liquid nitrogen, and stored at −80 °C until use.

### Cloning of piggyBac-PuroR-GADD34_Δ1-240/P2A/K3L

The PiggyBac transposon vector was constructed using PB-TRE3G (41) (Addgene #97421; gift from the Jennifer Doudna lab) as the backbone. The vector was linearized with NcoI and BsrGI restriction enzymes (New England Biolabs) according to the manufacturer’s instructions. The GADD34Δ1-240–P2A–K3L insert was amplified by PCR from the template plasmid pSBtet-Hygro-GADD34_Δ1-240/P2A/K3L (5) (Addgene #196136) using Q5 High-Fidelity DNA Polymerase and the following primers:

Forward: 5′-TCGTAAAGGTCTAGAGCTAGCCACCATGAAAGGAGCCAGGAAGACC-3′

Reverse: 5′-CTAACCGGTCAACCGGTCTTGTACATCACTGGTGGCGACACATAC-3′

Results of both reactions were verified by agarose gel electrophoresis. PCR fragment was treated with DpnI (NEB) overnight at 37 °C to remove template plasmids and purified using the DNA Clean & Concentrator-5 Kit (Zymo Research). Gibson Assembly was performed using the NEBuilder® HiFi DNA Assembly Master Mix (New England Biolabs) according to the manufacturer’s protocol. The assembled plasmid was sequence-verified by full plasmid sequencing and has been deposited to Addgene (Plasmid #249033).

### Human Primary Fibroblasts Cell Culture

WI-38 human primary fibroblasts (from UC Berkeley cell culture facility) were cultured in a 1:1 mixture of Ham’s F-12 and Dulbecco’s Modified Eagle Medium (DMEM; Gibco) supplemented with 10% Tet-system approved FBS (Gibco) and 1× penicillin-streptomycin (Gibco). Cells were maintained at 37 °C in a humidified atmosphere containing 5% CO₂ and routinely passaged at 70– 80% confluence using 0.05% trypsin-EDTA. All cell lines were confirmed to be mycoplasma-free by routine PCR-based testing.

### Generation of WI-38 fibroblasts with a stably integrated GADD34Δ1-240/K3L construct

A stable WI-38 human primary fibroblast line expressing the GADD34Δ1-240/K3L construct was generated using the PiggyBac transposon system (42). Cells were nucleofected with the 500 µg of pPB-TRE3G-GADD34Δ/K3L plasmid together with the 500 µg of Super PiggyBac transposase expression vector (a gift from the Jonathan Braverman lab) using the Amaxa 4D-Nucleofector X Unit (Lonza) and P3 kit for Primary Cell kit. After 48 h of recovery, transfected cells were selected with 0.5 µg/mL puromycin (Gibco) for 5–7 days. The efficiency of integration and selection was monitored by visual inspection of culture confluence and viability. Stably transfected WI-38 cells were expanded and subsequently used for functional assays.

### iPSC Cell Culture

Human induced pluripotent stem cells (iPSCs; KOLF2.1J line, UC Berkeley cell culture facility) were maintained under feeder-free conditions using mTeSR plus medium (STEMCELL Technologies) on tissue culture plates coated with growth factor–reduced Matrigel (Corning). For plate coating, Matrigel was diluted according to the manufacturer’s recommendation in cold DMEM/F-12 (Gibco) and incubated at 37 °C for at least 1 h before use. Cells were cultured at 37°C in a humidified incubator with 5% CO_2_, 5% O_2_ and passaged every 3–4 days upon reaching approximately 80–90% confluency. Routine passaging was performed using ReLeSR (STEMCELL Technologies) following the manufacturer’s instructions to detach colonies while preserving small clumps of undifferentiated cells. Detached cell aggregates were resuspended in fresh mTeSR plus medium and seeded onto newly Matrigel-coated plates at a 1:6–1:10 split ratio. For applications requiring single-cell suspensions, such as genome editing or directed differentiation, iPSCs were dissociated using Accutase (Gibco) for 10 min at 37 °C. The resulting single-cell suspensions were centrifuged at 200 × *g* for 3 min, resuspended in mTeSR plus medium containing 10 µM Y-27632, and replated at a density of 2-3 × 10^5^ cells/cm^2^. Cells were routinely monitored for colony morphology, spontaneous differentiation, and mycoplasma contamination (PCR-based testing). Only cultures displaying compact colonies with well-defined borders and a high nuclear-to-cytoplasmic ratio were used for downstream experiments.

### Mesoderm differentiation (pluripotency validation assay)

Human KOLF2.1J iPSCs were seeded on Matrigel-coated 6-well plates at ∼2.5 × 10^5^ cells per well in mTeSR Plus medium (STEMCELL Technologies) and allowed to attach overnight. Mesoderm differentiation was performed according to (43, 44) with minor modifications. Briefly, mesoderm induction was performed for 48 h using RPMI-1640 (Gibco) supplemented with Activin A (5 ng/mL; PeproTech), CHIR99021 (5 µM; TOCRIS), and BMP4 (5 ng/mL; PeproTech). The induction medium was prepared fresh and replaced once at 24 h. Cultures were maintained at 37 °C, 21% O_2_, 5% CO_2_. At 48 h, cells were fixed in 4% paraformaldehyde (15 min, RT), washed in PBS, and processed for immunofluorescence.

### Generation of human iPSCs with a stably integrated GADD34Δ1-240/K3L construct

Stable integration of the GADD34Δ1-240/K3L construct in human iPSCs was achieved using the PiggyBac transposon system described above. iPSCs were seeded onto Matrigel-coated plates (Corning) in mTeSR Plus medium and pre-treated with 10 µM Y-27632 ROCK inhibitor (TOCRIS) for 24 h prior to nucleofection to enhance cell survival. Cells were then nucleofected with the 500 ng of pPB-TRE3G-GADD34Δ/K3L plasmid together with the 500 ng of Super PiggyBac transposase expression vector (a gift from the Braverman lab) using the Amaxa 4D-Nucleofector X Unit (Lonza) and P3 kit for Primary Cell (Lonza) according to the manufacturer’s instructions. Following nucleofection, cells were allowed to recover for 48 h in mTeSR plus medium supplemented with 10 µM Y-27632, then subjected to puromycin selection (0.5 µg/mL) for 5–7 days. Surviving colonies were expanded on Matrigel-coated plates and maintained under standard feeder-free iPSC culture conditions.

### Human iPSC Culture and Differentiation into Cardiomyocytes

Human induced pluripotent stem cells (iPSCs) (KOLF2.1J line, UC Berkeley cell culture facility) were maintained in mTeSR plus medium (STEMCELL Technologies) on Matrigel-coated plates (Corning). Plates were coated by diluting Matrigel in DMEM/F12 (Gibco) and incubating at 37 °C for at least 1 h. Cells were passaged using Accutase (Gibco) and replated at a density of ∼0.25-0.5 × 10^6^ cells per well in 6-well plates in mTeSR plus supplemented with 10 µM ROCK inhibitor Y-27632 (STEMCELL Technologies). After 24 h, cultures were maintained at 37 °C in 5% CO_2_, with daily medium exchange until reaching >90% confluence.

Cardiomyocyte differentiation was performed following the protocol from (45–48) with minor modifications. Briefly, differentiation was initiated (Day 0) by replacing the maintenance medium with RPMI 1640 (Gibco) supplemented with B-27 Supplement minus insulin (Gibco) and 4 µM CHIR-99021 (TOCRIS). After 48 h (Day 2), medium was replaced with RPMI-B27 minus insulin. On Day 3, medium was exchanged for RPMI-B27 minus insulin containing 5 µM IWP-4 (TOCRIS), and cells were incubated for 48 h. On Day 5 and Day 7, cells were washed with PBS and fed with fresh RPMI-B27 minus insulin. On Day 9 and Day 11, media were switched to RPMI 1640 supplemented with B-27 with insulin (Gibco), at which point spontaneous contractile activity was typically observed by Day 10-11. For metabolic selection, media were replaced on Day 13 and Day 14 with DMEM without glucose (Gibco) supplemented with 4 mM sodium L-lactate (Sigma-Aldrich). Following selection, cultures were returned to RPMI-B27 with insulin on Day 15 and maintained until replating.

Following differentiation and lactate-based metabolic selection, hiPSC-derived cardiomyocytes (hiPSC-CMs) were replated and expanded according to the protocol (49, 50). Briefly, hiPSC-CMs were enzymatically dissociated using pure TrypLE Select (10×) (Gibco) at 37 °C for ∼15 min until cells detached completely. The detached cells were collected in pre-warmed RPMI 1640 medium, centrifuged (200 × *g*, 3 min), and resuspended in pre-heated cardiomyocyte replating medium (RPMI + B-27 + 10% KO serum replacement (Gibco) + 10 µM ROCK inhibitor Y-27632 (STEMCELL Technologies) + 2 µM CHIR-99021 (TOCRIS)). Cells were counted and seeded on freshly Matrigel-coated culture flasks at a split ratio of 1:10–1:20 (approximately 3 × 10^4^ cells/cm^2^) and incubated at 37 °C, 5% CO_2_, 21% O_2_, and 90% humidity. After 24 h of recovery, replating medium was replaced with cardiomyocyte expansion medium (RPMI + B-27 + 2 µM CHIR-99021 (TOCRIS)). Media were refreshed every 2–3 days, and cultures were passaged upon reaching 70–80% confluency. hiPSC-CMs were expanded through up to three passages (P1–P3). Each passage approximately doubled the cell number, enabling large-scale expansion while preserving cardiomyocyte identity.

### Immunocytochemistry

Cells were washed once with phosphate-buffered saline (PBS, pH 7.4, Gibco), then fixed with 3% paraformaldehyde (PFA) (Electron Microscopy Sciences) in PBS at room temperature for 15 min. After two additional PBS washes (5 min each), samples for intracellular markers were permeabilized in 0.3% Triton-X100 (Sigma) in PBS for 30 min at room temperature. Non-specific binding was blocked by incubating in 3% bovine serum albumin (BSA) in PBS for 1 h at room temperature. Samples were incubated overnight at 4 °C with primary antibodies (anti-OCT4 (sc-5279), anti-SSEA4 (sc-21704), anti-Actinin (sc-17829), anti-Troponin (sc-20025), anti-Brachyury (A-4) (sc-374321), all from Santa Cruz Biotechnology) diluted in blocking solution. The following day, cells were washed with PBS and incubated with appropriate secondary antibodies (anti-mouse IgG (H+L) Alexa Fluore 488 donkey secondary antibody from Invitrogen, A21202; or anti-mouse IgG (H+L) Alexa Fluore 594 donkey secondary antibody from Invitrogen, A21203) for 1 h at room temperature, followed by a final PBS wash and nuclear counterstaining with 0.1 μg/mL DAPI (Thermo Fisher Scienic). Stained cultures were imaged using a fluorescence or bright-field microscope to confirm high expression of cell markers, evaluate cellular morphology, and quantify marker-positive cells or colonies as required.

### Cytochemical Alkaline Phosphatase Staining

Fixed iPSCs were equilibrated in 100 mM Tris-HCl (pH 9.5) for 10 min and subsequently incubated with NBT/BCIP substrate solution (Roche) at room temperature for 2 h. This reaction produced a blue-purple precipitate marking alkaline phosphatase–positive colonies. Stained colonies were imaged directly on the culture plate using an iPhone 11 under bright-field illumination.

### Preparation of WI-38 human primary fibroblast and cardiomyocyte cell extracts

First, approximately 1.2-1.5 million cells were seeded per 150 mm plate in cell-specific media, described above. The next day, the expression of GADD34Δ and K3L was induced by adding 1 µg/mL of doxycycline (Takara). After one additional day, cells were collected by scraping, washed with ice-cold DPBS (Gibco), and suspended with an equal volume of lysis buffer (10 mM HEPES pH 7.4, 10 mM KOAc, 0.5 mM Mg(OAc)_2_, and 5 mM DTT). After incubation on ice for 45 min, cells were lysed by pushing through a 1-ml syringe with a 26G needle about 15 times, followed by centrifugation at 15000 x *g* for 1 min. After centrifugation, the supernatant was aliquoted to avoid freeze-thaw cycles and flash-frozen in liquid nitrogen.

### *In vitro* transcription reactions

*In vitro* transcription reactions were performed using PCR products generated with primers encoding a flanking T7 RNA polymerase promoter and a poly-A tail. Reactions were set up as previously described (51), with 20 mM Tris-HCl pH 7.5, 35 mM MgCl_2_, 2 mM spermidine, 10 mM DTT, 1 u/mL pyrophosphatase (Sigma), 7.5 mM of each NTP, 0.2 u/mL RiboLock RNase Inhibitor (ThermoFisher), 0.1 mg/mL T7 RNA polymerase and 40 ng/μL PCR-generated DNA.

After 3 h incubation at 37 °C, 0.1 u/μL DNase I (Promega) was added to the reactions, which were incubated at 37 °C for 30 min to remove the template DNA. RNA was precipitated for 2–3 h at −20 °C after adding 0.5x volume of 7.5 M LiCl/50 mM EDTA, and the resulting pellet was washed with cold 70% ethanol and dissolved with RNase-free water. The mRNA was further purified by using a Zymo RNA Clean and Concentrator (Zymo Research) before use in *in vitro* translation reactions.

### *In vitro* translation reactions

Translation reactions were set up according to a previously published procedure (5, 10) with modifications. For a 10 μL mRNA-dependent reaction, 5 μL of cell extract was used in a buffer containing final concentrations of 52 mM HEPES pH 7.4 (Takara), 35 mM KGlu (Sigma), 1.75 mM Mg(OAc)_2_ (Invitrogen), 0.55 mM spermidine (Sigma), 1.5% Glycerol (Fisher Scientific), 0.7 mM putrescine (Sigma), 5 mM DTT (Thermo Scientific), 1.25 mM ATP (Thermo Fisher Scientific), 0.12 mM GTP (Thermo Fisher Scientific), 100 µM L-Arg; 67 µM each of L-Gln, L-Ile, L-Leu, L-Lys, L-Thr, L-Val; 33 µM each of L-Ala, L-Asp, L-Asn, L-Glu, Gly, L-His, L-Phe, L-Pro, L-Ser, L-Tyr; 17 µM each of L-Cys, L-Met; 8 µM L-Trp, 20 mM creatine phosphate (Roche), 60 µg/mL creatine kinase (Roche), 4.65 µg/mL myokinase (Sigma), 0.48 µg/mL nucleoside-diphosphate kinase (Sigma), 0.3 u/mL inorganic pyrophosphatase (Thermo Fisher Scientific), 100 µg/mL total calf tRNA (Sigma), 0.8 u/μL RiboLock RNase inhibitor (Thermo Scientific) and 1000 ng mRNA. The optimal concentration of the magnesium and potassium ions was determined for each new preparation of cellular extract, with the conditions above representative of typical final conditions. Reactions were incubated for 60 min at 32 °C, and nanoluciferase activity was monitored using the Nano-Glo Luciferase Assay Kit (Promega) in a Microplate Luminometer (Veritas). In experiments testing cap-dependent translation with the m^7^G-capped NanoLuciferase reporter with an *HBB* 5’-UTR, 250 µM m^7^GTP (Santa Cruz Biotechnology) was added to the reactions. To account for batch-to-batch variability, all assays presented in a given figure were carried out using the same *in vitro* translation extract and same preparation of mRNAs.

For the *in vitro* translation of GFP experiments, coupled transcription-translation reactions based on the HeLa and HEK293T extracts were supplemented with 100 ng/μL EMCV-GFP mRNA transcribed from pCFE1-GFP plasmid (Thermo Scientific). After three hours incubation, reactions were transferred directly into a black 384-well plate with clear bottom (Greiner). The GFP fluorescent signal was measured using a Tecan SPARK plate reader at ex/em: 482/512 nm.

### Western blot analysis

Samples were boiled in Bolt LDS Sample loading buffer (Thermo Fisher Scientific) containing Bolt/NuPAGE reduction buffer (Thermo Fisher Scientific) at 70 °C for 10 min. Samples were resolved on 4%–12% Bolt Bis-Tris Plus protein gels (Thermo Fisher Scientific) using 1x Bolt MES or MOPS SDS running buffer (Thermo Fisher Scientific) containing 1x NuPAGE Antioxidant (Thermo Fisher Scientific) according to the manufacturer’s instructions. Gels were transferred to nitrocellulose membranes using a Power Blot system (Thermo Fisher Scientific) using medium-range manufacturer parameters. Membranes were blocked with 5% nonfat dry milk (Bioworld) in PBST (10 mM Tris-HCl, pH 8 (Invitrogen), 1 mM EDTA (Invitrogen), 0.1% Triton X-100 (Sigma), 150 mM sodium chloride (Sigma)) for 1 hr at room temperature (RT) with gentle rocking. Blots were washed in PBST 3 times and then incubated with the indicated primary antibodies overnight at 4 °C with gentle rocking. Blots were washed with PBST 3-4 times over 45-60 min at RT with gentle rocking, then incubated with secondary antibodies diluted in 5% milk in PBST for 1 hr with gentle rocking at RT. Membranes were washed again with PBST, 3-4 times over 45-60 min at RT, developed using SuperSignal West Pico Plus ECL substrate (Thermo Fisher Scientific) and SuperSignal West Femto Maximum Sensitivity Substrate (Thermo Fisher Scientific) if needed and imaged on an Ibright CL1000 (Thermo Fisher Scientific) system. Results shown are representative of at least two independent experiments.

### Phos-tag gels

For Phos-tag gel immunoblotting (52), samples were resolved on homemade 12 % Bis-Tris, pH 6.8, SDS-PAGE gels containing 50 μM Phos-tag (Wako, AAL-107) and 100 μM ZnCl_2_. Samples were boiled in Bolt LDS Sample loading buffer (Thermo Fisher Scientific) containing Bolt/NuPAGE reduction buffer (Thermo Fisher Scientific) at 70 °C for 10 min. Samples were resolved using 1x Bolt MES SDS running buffer (Thermo Fisher Scientific) containing 1x NuPAGE Antioxidant (Thermo Fisher Scientific) according to the manufacturer’s instructions. Gels were run using a constant 90 V until the bromophenol blue dye reached the bottom buffer. Before transfer to the nitrocellulose membrane, gels were soaked in 1 mM EDTA for 10 min with agitation to remove the Zn^2+^ ions. Gels were transferred to nitrocellulose membranes using a Power Blot system (Thermo Fisher Scientific) using medium-range manufacturer parameters. Membranes were blocked with 5% nonfat dry milk (Bioworld) in PBST (10 mM Tris-HCl, pH 8 (Invitrogen), 1 mM EDTA (Invitrogen), 0.1% Triton X-100 (Sigma), 150 mM sodium chloride (Sigma)) for 1 hr at RT with gentle rocking. Blots were washed in PBST 3 times and then incubated with the indicated primary antibodies overnight at 4 °C with gentle rocking. Blots were washed with PBST 3-4 times over 45-60 min at RT with gentle rocking, then incubated with secondary antibodies diluted in 5% milk in PBST for 1 hr with gentle rocking at RT. Membranes were washed again with PBST 3-4 times over 45-60 min at RT, developed using SuperSignal West Pico Plus ECL substrate (Thermo Fisher Scientific) and SuperSignal West Femto Maximum Sensitivity Substrate (Thermo Fisher Scientific) (if needed) and imaged on an Ibright CL1000 (Thermo Fisher Scientific) system. Results shown are representative of at least two independent experiments. The percents of the phosphorylation were estimated by comparing the intensities of the bands of phosphorylated and not phosphorylated forms of factors using ImageJ software (53). The background in the gel images was estimated using a rectangular region with equal dimensions positioned immediately above the band corresponding to the phosphorylated form of the proteins. A similar rectangular region was also selected below the non-phosphorylated band.

### Antibodies

The following antibodies were used in this study. Antibodies for eIF2*a* (9722S), PKR D7F7 (12297T), phospho-eIF2*a* (S51) (9721S), eEF2 (2332S) and phospho-eEF2 (T56) (2331S) were from Cell Signaling Technology. Antibodies for RPS19 (A304-002A) and eEF2K (A301-686A-T) were purchased from Bethyl Laboratories Inc. Anti-mouse IgG-HRP (sc-525409) and GADD34 (sc-373815) were from Santa Cruz Biotechnology. Anti-rabbit ECL IgG-HRP (NA934V) was from Thermo Fisher Scientific.

## Results

### Blocking eIF2α Phosphorylation Substantially Improves Translation in Expi293F Human Cell Extracts

Building on our prior work establishing human cell–derived translation systems (5), we first determined how specific phosphorylation events influence translational output in human cell-free translation systems derived from Expi293F suspension cells because they grow to very high densities, and enable the efficient generation of large quantities of biomass suitable for extract preparation (**Fig. 1D**). We introduced targeted mutations in *EIF2S1* (eIF2α) and *EEF2* (eEF2), designed to eliminate inhibitory phosphorylation at Ser52 in eIF2α and Thr57 in eEF2 (**Fig. 1E**). To introduce precise point mutations into essential translation factors without generating double-strand breaks, we employed prime editing (38, 54–57) (**Fig. S1A**), a versatile genome-editing technology that couples a catalytically impaired Cas9 nickase to a reverse transcriptase (RT). Using an engineered prime-editing guide RNA (pegRNA) (**Fig. S1B**), the Cas9–RT complex nicks the target DNA strand and uses the pegRNA-encoded reverse-transcription template to directly write the desired sequence into the genome (**Fig. S1A**). The pegRNA also contains a primer-binding site (PBS) that enables the reverse transcriptase to initiate DNA synthesis from the nicked strand, ensuring high-fidelity incorporation of the edit (**Fig. S1B**). To enhance efficiency and reduce byproducts, we used engineered pegRNAs (epegRNAs) with structured 3′ extensions that protect against exonuclease degradation (**Fig. S1B**). A second short guide RNA (nicking sgRNA) was co-delivered to introduce a nick on the non-edited DNA strand, biasing repair toward the newly edited sequence and improving conversion efficiency (**Fig. S1A**).

The prime editing construct (PEmax) (38, 56)—encoding Cas9(H840A)-RT—was cloned into a Sleeping Beauty (SB) transposon (37) vector under a tetracycline-inducible promoter (**Fig. S1C**). This vector was first stably integrated into Expi293F suspension cells using the SB transposase (**Fig. S1D, upper panel**). Following puromycin selection to enrich for successfully transposed cells, we introduced plasmids encoding the appropriate epegRNA and nicking sgRNA by nucleofection (**Fig. S1D, lower panel**). This two-step workflow decoupled stable installation of the editing machinery from transient guide delivery, allowing controlled induction of PEmax with doxycycline to minimize cellular toxicity, increase prime editing efficacy and reuse of the same parental line for additional edits. We applied this approach to introduce the Ser52 to Ala substitution in *EIF2S1* (encoding eIF2α) and the Thr57 to Ala substitution in *EEF2* (encoding eEF2) (**Fig. 1E**). For each locus, epegRNAs with variable primer-binding site (PBS) and reverse-transcription template (RTT) lengths were screened to optimize editing efficiency (**Fig. S2A-B and Fig. S3A-B**). Among *EIF2S1* designs, an epegRNA with a 13-nt PBS and 24-nt RTT yielded the highest editing frequency, with minimal indel formation (<3%), and the “–54” sgRNA position produced the most accurate edits (**Fig. S2C**). Three independent *EIF2S1-S52A* clones were isolated and confirmed by sequencing and while exhibiting normal proliferation (**Fig. S2D**). A parallel workflow for *EEF2* produced verified *EEF2-T57A* clones (**Fig. S3C–E**).

Genome-edited Expi293F cells were cultured in suspension, harvested, and lysed under identical conditions to generate extracts for *in vitro* translation (IVT) reactions programmed with a Nanoluciferase (NanoLuc) reporter mRNA (**Fig. 1D**) (58). The translation assays (**Fig. 1F**) demonstrated that only the eIF2α-S52A mutation significantly increased translational yield—by approximately 30-fold compared with wild-type extracts—indicating that inhibition of initiation through eIF2α phosphorylation constitutes the dominant bottleneck in Expi293F lysates. To ensure that enhanced translation in eIF2α S52A lysates was not specific to the NanoLuc reporter or EMCV IRES context, we also tested alternative templates encoding *NanoLuc* under the HCV IRES and *copGFP* under the EMCV IRES, both of which showed similarly elevated translation (**Fig. S4A, B**). Consistent with these findings, eIF2α S52A extracts also supported robust translation of the reporter mRNA with m^7^G-capped *HBB* (human β-globin) 5’UTR in a cap-dependent manner, further confirming that the enhancement extends beyond IRES-driven reporters (**Fig. S4C**). By contrast, eliminating eEF2 phosphorylation by T57A genome edits only improved translation activity of the cell extract by ∼50% (**Fig. 1F**). Additionally, by using CRISPR–Cas9 targeting of exon 7 (**Fig. S5A–B**) we generated eEF2 kinase knockout clones, which eliminated eEF2K protein expression without affecting cell growth (**Fig. S5C**). Knockout of the eEF2K did not result in substantially increased translation efficiency of the cell extract, although it completely abolished eEF2 phosphorylation (**Fig. S6**), further indicating that elongation control contributes minimally to steady-state translation limitation in cell-free conditions. Together, these data demonstrate that phosphorylation of eIF2α, but not eEF2, is the primary molecular constraint suppressing protein synthesis in Expi293F human cell lysates.

### Partial Relief of Translation Inhibition by PKR Knockout

To determine whether a kinase responsible for eIF2α phosphorylation during extract preparation could alternatively be targeted to improve lysate productivity, we generated PKR knockout (PKR⁻/⁻) Expi293F cells (**Fig. 2A**). The double-stranded RNA–activated protein kinase (PKR; encoded by *EIF2AK2*) is one of the four known eIF2α kinases that phosphorylate Ser52 in response to viral or stress signals, and previous biochemical analysis of human translation extracts implicated PKR activation during cell lysis as a major driver of inhibitory eIF2α phosphorylation (12, 59). Using a CRISPR–Cas9–based approach, we disrupted *EIF2AK2* by introducing a small frameshift deletion within exon 15 (**Fig. S7A**). Immunoblotting confirmed complete loss of PKR protein in three independent clones (**Fig. S7B**), all of which proliferated similarly to wild-type Expi293F cells (**Fig. S7C**). Extracts prepared from these PKR⁻/⁻ lines exhibited a clear increase in translation activity relative to wild-type lysates but did not reach the high yields observed for the eIF2α-S52A mutant (**Fig. 2B**). Western blot analysis of eIF2α phosphorylation in the corresponding lysates (**Fig. 2C**) revealed that wild-type extracts displayed nearly complete phosphorylation of eIF2α (∼100%), whereas phosphorylation was fully abolished in the eIF2α S52A mutant. In contrast, the PKR-deficient extracts retained ∼30% residual phosphorylation, consistent with only partial relief of the initiation block. These data establish PKR as the primary but not exclusive source of inhibitory eIF2α phosphorylation in Expi293F lysates. Eliminating PKR improves translational capacity by mitigating stress-induced phosphorylation but cannot fully replicate the high activity achieved by direct mutation of the phosphorylation site. Thus, while PKR knockout provides a viable engineering route when genome editing of *EIF2S1* is not feasible, complete removal of the eIF2α regulatory site remains the most effective strategy for generating high-yield human cell-free translation systems.

**Figure 2.**
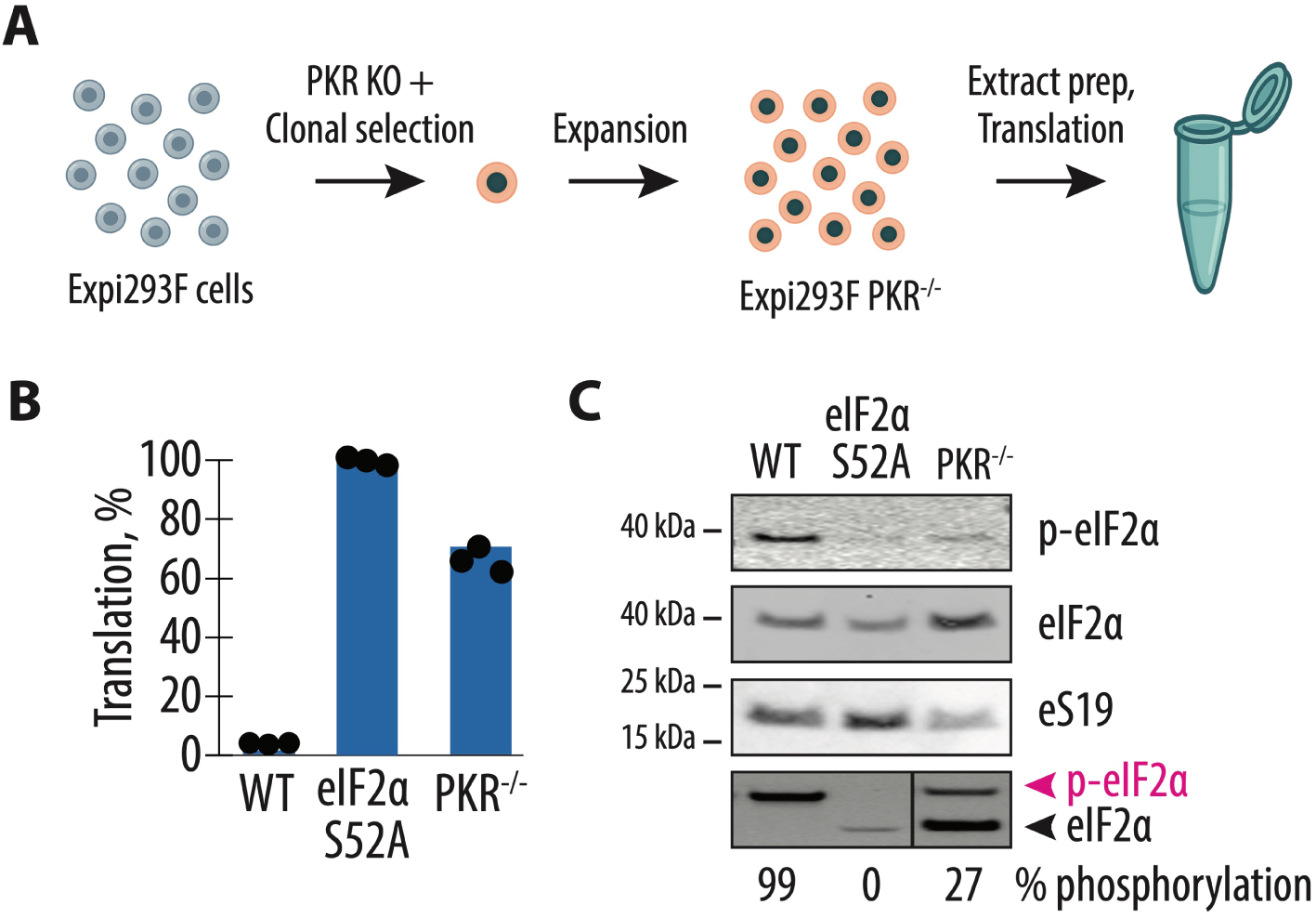
PKR knockout partially relieves the eIF2α phosphorylation block. (**A**) Expi293F cells were edited to generate PKR⁻/⁻ clones (*EIF2AK2* knockout) that were expanded, and cell extract was prepared for IVT assays. (**B**) NanoLuc IVT readout (% of eIF2α-S52A set to 100%). WT extracts show low activity; eIF2α-S52A shows high activity; PKR⁻/⁻ shows an intermediate increase. Each point is a biological replicate. All experiments were performed in biological triplicates. (**C**) Phos-tag SDS-PAGE and corresponding immunoblot of the same lysates probed for phospho-eIF2α (Ser52) and total eIF2α. The phos-tag gel separates phosphorylated and non-phosphorylated forms of eIF2α, allowing precise quantification of phosphorylation stoichiometry shown below. WT extracts display nearly complete phosphorylation (∼100%), eIF2α-S52A lysates lack any detectable phospho-signal (∼0%), and PKR⁻/⁻ extracts retain only ∼30% residual phosphorylation, indicating partial relief of initiation inhibition. Gels are representative of two independent experiments.

### Inducible Co-expression of GADD34 and K3L Restores Translation Efficiency of Primary Human Fibroblasts cell extracts

While Expi293F-based cell-free translation systems offer high translational capacity and scalability for recombinant protein synthesis, they rely on an immortalized cell line, which may not fully recapitulate physiological translation regulation. To expand this platform beyond industrial cell types, we sought to establish a human cell–derived *in vitro* translation system based on primary cells, which could better model endogenous regulatory environments. As a representative model of normal human primary cells, we selected WI-38 fibroblasts, which retain intact stress-response pathways and non-transformed physiology. Because precise genome editing approaches such as prime editing are inefficient and technically challenging in primary cells, we instead implemented an expression-based rescue strategy to alleviate inhibitory eIF2α phosphorylation and restore translational activity. To achieve this, we stably introduced a bicistronic GADD34Δ/K3L construct into WI-38 fibroblasts using the piggyBac transposon system (GADD34Δ lacks the amino-terminal 240 amino acids, and retains full activity (35)) (**Fig. 3A**). GADD34 recruits PP1 to catalyze eIF2α dephosphorylation, while K3L functions as a viral decoy substrate that sequesters stress-activated kinases, preventing further phosphorylation. Unlike the Sleeping Beauty vector previously used for prime editing in Expi293F cells, the piggyBac platform provides superior transposition efficiency in non-dividing or slowly dividing cells, making it particularly suitable for primary cultures.

**Figure 3.**
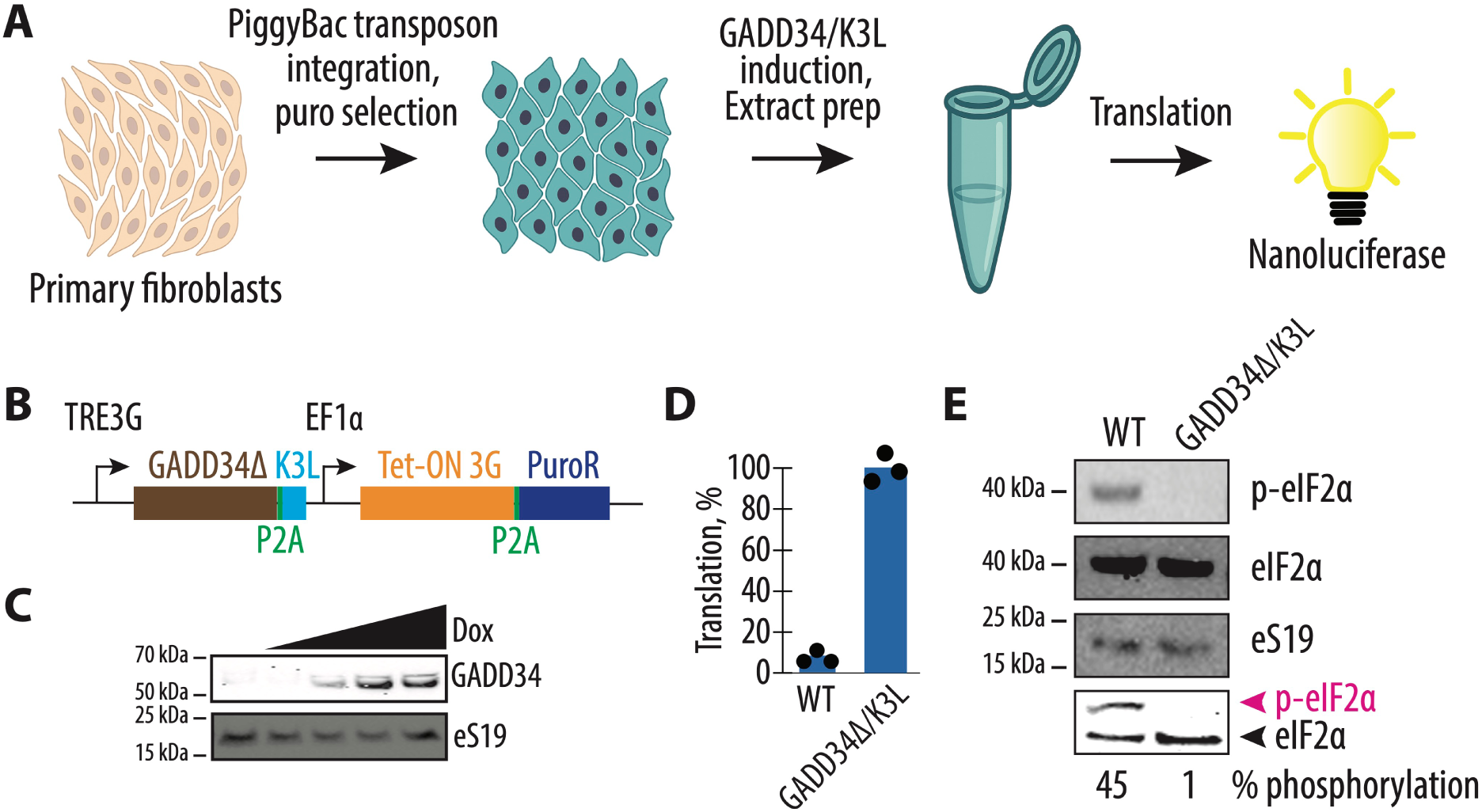
Inducible GADD34Δ/K3L co-expression suppresses eIF2α phosphorylation and restores translation efficiency in primary human fibroblast extracts. **(A)** Workflow for establishing translation-competent primary human fibroblasts. WI-38 primary fibroblasts were transfected with a piggyBac transposon carrying the inducible GADD34Δ/K3L cassette, selected with puromycin, and induced with doxycycline prior to extract preparation and *in vitro* translation (IVT) analysis. **(B)** Design of the inducible bicistronic expression construct. The TRE3G promoter drives expression of GADD34Δ-P2A-K3L, while the EF1α promoter constitutively expresses Tet-ON 3G and a puromycin-resistance cassette (PuroR) separated by a P2A sequence. (**C**) Immunoblot analysis of doxycycline-inducible GADD34Δ expression. Increasing doxycycline concentrations (0–1 µg/mL) induced dose-dependent accumulation of GADD34Δ protein. Ribosomal protein eS19 serves as loading control. Representative of two independent experiments. **(D)** *In vitro* translation activity of wild-type and GADD34Δ/K3L-expressing fibroblast extracts programmed with NanoLuc mRNA. Induced GADD34Δ/K3L expression results in an increase in translation yield compared with wild-type extracts. Each point represents a biological replicate; bars show mean. All experiments were performed in biological triplicates. **(E)** Immunoblot analysis of the same lysates probed for phospho-eIF2α (Ser52), total eIF2α, and ribosomal protein eS19 (loading control). On the bottom, phosphorylated and non-phosphorylated eIF2α forms were resolved by phostag SDS-PAGE, and phosphorylation stoichiometry was quantified from band intensities (values below lanes). Gels are representative of two independent experiments.

We previously developed the GADD34Δ /K3L co-expression approach (5) to counteract eIF2α phosphorylation in mammalian translation extracts. Here, we adapted it into a modern, inducible expression format, placing the transgene under control of the TRE3G promoter and Tet-ON 3G regulatory system (**Fig. 3B**). This design allows doxycycline-inducible expression of GADD34Δ and K3L with minimal basal activity and is compatible with a wide range of mammalian cell types, including primary and patient-derived cells. Upon doxycycline induction, fibroblast lysates expressing GADD34Δ/K3L showed a dramatic increase in translation efficiency compared to wild-type extracts (**Fig. 3C**). Western blotting confirmed that phospho-eIF2α levels decreased from ∼45% in wild-type to nearly undetectable levels in GADD34Δ/K3L-expressing cells (**Fig. 3D**). Both IRES-driven and cap-dependent reporters translated efficiently in the engineered fibroblast extracts (**Fig. S8**). These results demonstrate that controlled co-expression of GADD34Δ and K3L effectively eliminates the inhibitory eIF2α phosphorylation and restores robust translation in non-immortalized human cells. This expression-based rescue strategy provides a practical alternative to genome editing, enabling generation of translationally active lysates from cell types where precise genetic modification is not feasible.

### Inducible GADD34Δ/K3L Expression Restores Translation Efficiency in iPSC-Derived Cardiomyocyte Lysates

We next sought to determine whether similar principles could be extended to human induced pluripotent stem cells (iPSCs). Human iPSCs are a particularly attractive platform, as they can be differentiated into virtually any somatic lineage, enabling cell-type–specific investigation of human translation under near-physiological conditions. Among these, KOLF2.1J iPSCs are a well-defined, genetically stable line extensively used in large-scale differentiation pipelines and disease-modeling consortia. We first attempted to use the prime editing approach devised for Expi293F cells to make the Ser52Ala mutation and block eIF2α phosphorylation. In our hands, we were unable to stably introduce the S52A mutation in *EIF2S1* in iPSCs, possibly due to the essential role this phosphorylation plays in stem cells (15, 60, 61). Because direct genome editing of eIF2α in iPSCs is seemingly incompatible with iPSC propagation and subsequent differentiation efficiency, we turned to an expression-based approach to overcome translational suppression. Specifically, we implemented the doxycycline-inducible bicistronic cassette encoding GADD34Δ and K3L described above to reverse or bypass eIF2α phosphorylation (**Fig. 4A**). The construct was stably integrated into the iPSC genome using the piggyBac transposon system (**Fig. 3B**), which provides superior transposition efficiency and long-term expression stability in slowly dividing or post-mitotic cells compared to Sleeping Beauty used for Expi293F genome editing.

**Figure 4.**
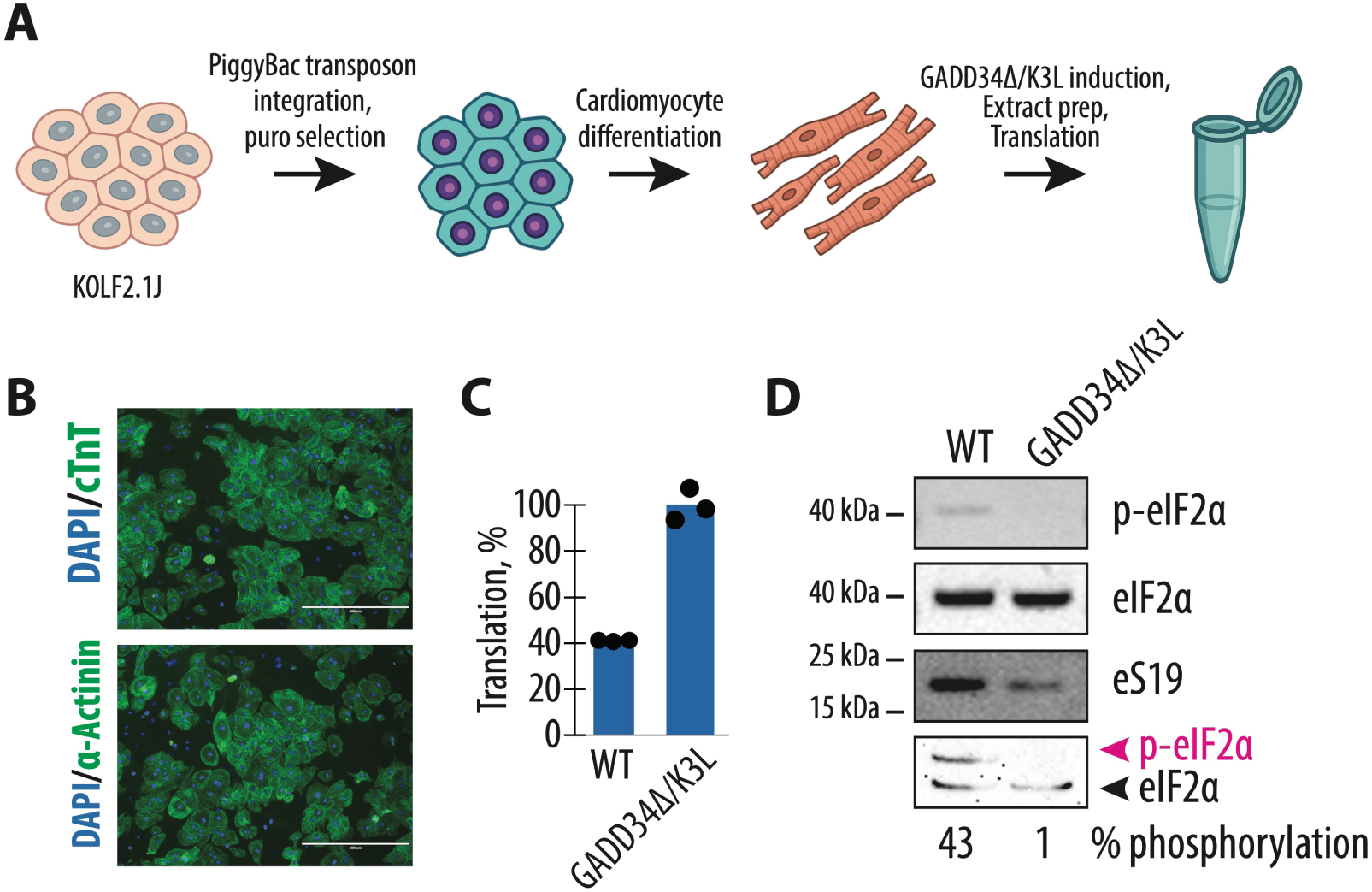
Inducible GADD34Δ/K3L expression restores translational activity in iPSC-derived cardiomyocytes. **(A)** Schematic overview of the experimental workflow. Human KOLF2.1J induced pluripotent stem cells (iPSCs) were stably transposed with a piggyBac vector encoding a doxycycline-inducible bicistronic GADD34Δ/K3L cassette and selected with puromycin. Following clone expansion, engineered iPSCs were differentiated into cardiomyocytes using a small-molecule CHIR99021/IWP-2 protocol. After metabolic selection, differentiated cells were induced with doxycycline to express GADD34Δ/K3L, lysed under standardized conditions, and used to prepare translation-competent extracts for *in vitro* translation (IVT) assays. (**B**) Immunofluorescence validation of cardiomyocyte identity. Representative fluorescence micrographs show cells stained for cardiac troponin T (cTnT) and α-actinin (green) with DAPI (blue) counterstain. Differentiation efficiency exceeded 90 %, and cells displayed organized sarcomeric structures typical of functional cardiomyocytes. Scale bars = 400 µm. (**C**) Quantification of translational activity in IVT assays programmed with capped NanoLuc reporter mRNA. Each point represents an independent biological replicate. All experiments were performed in biological triplicates. (**D**) Immunoblot analysis of eIF2α phosphorylation in WT and GADD34/K3L-induced cardiomyocyte lysates. Immunoblots were probed with antibodies against phospho-eIF2α (Ser52), total eIF2α, and ribosomal protein eS19 as a loading control. Data are representative of at least three independent extract-preparation experiments. Gels are representative of two independent experiments.

Following puromycin selection of transposed clones, GADD34Δ/K3L iPSCs were differentiated into cardiomyocytes using the chemically defined CHIR99021–IWP-2 protocol (**Fig. S9A**) (45–48). During differentiation, iPSCs were transitioned from mTeSR Plus to RPMI 1640 + B27 without insulin, sequentially exposed to Wnt activator (CHIR99021) and inhibitor (IWP-2), and subsequently matured in lactate-based medium for metabolic selection. This procedure consistently yielded highly pure cardiomyocyte populations, with differentiation efficiencies exceeding 90 %, as verified by immunofluorescence staining for cardiac troponin T (cTnT) and α-actinin (**Fig. 4B**). The resulting monolayers displayed organized sarcomeric structures and rhythmic contractions characteristic of functional cardiomyocytes. To evaluate the inducible system, cells were treated with increasing concentrations of doxycycline to activate the TRE3G promoter. Western blotting confirmed dose-dependent accumulation of GADD34 protein in cardiomyocytes (**Fig. S9B**), demonstrating robust inducibility of the transgene in this post-mitotic human lineage. Following induction, cardiomyocytes were lysed under standardized conditions to prepare translation-competent extracts. *In vitro* translation assays using NanoLuc reporter mRNA revealed that GADD34/K3L expression approximately doubled translational output relative to uninduced wild-type (WT) cardiomyocyte lysates (**Fig. 4C**). To determine whether this increase correlated with alleviation of the initiation block, we examined eIF2α phosphorylation status by immunoblotting and Phos-tag analysis. In WT cardiomyocyte lysates, roughly 40 % of eIF2α was phosphorylated, indicating substantial induction of the stress-responsive inhibitory mark prior to or during extract preparation. In contrast, doxycycline-induced GADD34/K3L expression reduced phosphorylation to near-undetectable levels, effectively restoring the pool of active eIF2α available for ternary-complex formation (**Fig. 4D**).

## Discussion

Here we identified inhibitory phosphorylation of eIF2α at Ser52 as the principal biochemical bottleneck limiting protein synthesis in human cell–derived *in vitro* translation (IVT) systems. By using prime editing, kinase knockout, and inducible dephosphorylation strategies to prevent eIF2α phosphorylation, we provide complementary genetic solutions that markedly enhance translational efficiency across multiple human cell types (**Fig. 5**). Together, these results establish a generalizable framework for constructing next-generation mammalian IVT platforms with improved yield, stability, and modularity.

**Figure 5.**
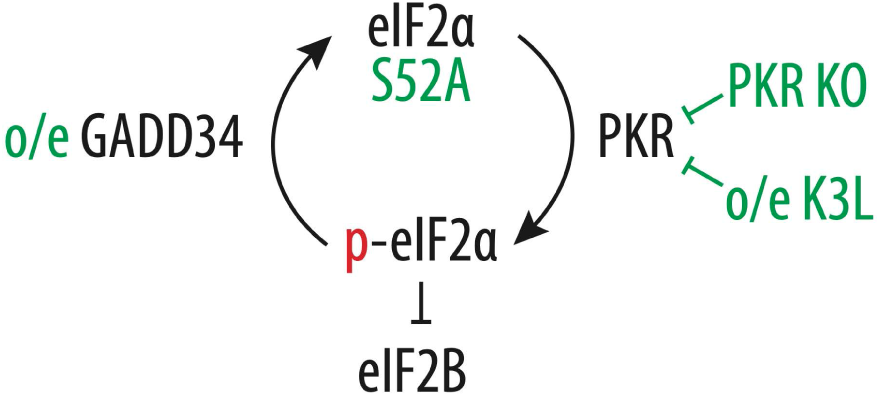
Genetic strategies to suppress eIF2α phosphorylation and restore translation initiation. Schematic summary of the regulatory network controlling eIF2α phosphorylation and the corresponding genetic interventions used in this study. Under stress or during extract preparation, PKR phosphorylates eIF2α on Ser52, converting it into p-eIF2α, which inhibits eIF2B and blocks GDP–GTP exchange required for ternary complex formation and translation initiation. Three distinct genetic approaches relieve this inhibition: (i) An eIF2α-S52A mutation prevents phosphorylation entirely; (ii) PKR knockout (PKR KO) or overexpression of the vaccinia virus K3L decoy (o/e K3L) inhibits PKR activity; and (iii) overexpression of GADD34Δ (o/e GADD34) promotes dephosphorylation of eIF2α. Together, these strategies converge to maintain eIF2α in a non-phosphorylated, translation-competent state, thereby enhancing protein synthesis in human cell-derived extracts.

Our systematic comparison of eIF2α and eEF2 modifications revealed that inhibitory phosphorylation of eIF2α, rather than eEF2, constitutes the dominant translational brake in Expi293F human cell extracts. Editing the *EIF2S1* locus to replace Ser52 with alanine completely abolished eIF2α phosphorylation and increased translational yield by nearly 30-fold compared with wild-type extracts (**Fig. 1F**). In contrast, mutation of eEF2 Thr57 or deletion of its kinase eEF2K did not improve translation efficiency, even though eEF2 phosphorylation was eliminated (**Fig. S6**). This indicates that under steady-state conditions, initiation rather than elongation remains rate-limiting in mammalian extracts—consistent with cellular models where eIF2α phosphorylation serves as a master switch of the integrated stress response (13, 14, 17).

Precise genetic elimination of the Ser52 phosphorylation site via prime editing provides a direct, stable, and effective means of improving extract performance without perturbing cell viability or requiring small-molecule additives. Because Expi293F cells are widely used for high-density suspension culture and recombinant-protein production, the engineered eIF2α-S52A Expi293F line represents a powerful chassis for industrial-scale lysate manufacturing and synthetic-biology applications (**Fig. 1E–F**). This single-point mutation combines the robustness of an established production host with enhanced translational capacity, establishing a foundation for reproducible, scalable, high-yield human IVT systems. The partial rescue observed in PKR-knockout Expi293F cells (**Fig. 2A–B**) underscores the redundancy among eIF2α kinases. Even in the absence of PKR, residual phosphorylation (∼30%) persisted, possibly mediated by GCN2 whose activation can be triggered by increased translational throughput and transient depletion of charged tRNAs. As translation accelerates in the absence of PKR, ribosome collision or imbalanced amino-acid usage may create stress that reactivates GCN2, maintaining a residual initiation block. This residual modification most likely reflects secondary activation of stress-responsive kinases, a mechanism we previously proposed (5). Thus, complete and stable elimination of eIF2α phosphorylation appears to be the most robust route to achieving high translational capacity in human systems.

While genome editing provided an efficient solution in Expi293F cells, prime editing and Cas9 dsDNA break repair-mediated editing remain challenging in non-dividing or primary human cells. To suppress eIF2α phosphorylation in more physiologically relevant cell types, we implemented an inducible GADD34Δ/K3L expression system in primary fibroblasts and iPSC-derived cardiomyocytes (**Figs. 3–4**). This bicistronic design, delivered via PiggyBac transposition (42) and controlled by a TRE3G/Tet-ON 3G promoter, enabled tight and reversible expression even in post-mitotic cells. Upon induction, GADD34Δ/K3L expression completely eliminated eIF2α phosphorylation and restored translation. These findings establish that transient induction of GADD34/K3L efficiently relieves translational repression in differentiated human cell types, recapitulating the beneficial effects of genome editing. Importantly, this result demonstrates that the biochemical bottleneck observed in immortalized lines—stress-activated phosphorylation of eIF2α—also constrains protein synthesis in primary cells. By providing a modular, reversible tool for translational activation, the inducible system enables production of high-activity lysates from editing-refractory human cells such as primary fibroblasts and iPSC-derived cardiomyocytes. Together, these data reinforce the universality of eIF2α phosphorylation as a central inhibitory mechanism and highlight the broad applicability of expression-based rescue strategies across cellular contexts.

A key advantage of the inducible GADD34/K3L system is its compatibility with developmental and differentiation workflows. Expression can be activated only after the completion of cardiomyocyte differentiation, preventing interference with the signaling networks that guide iPSC maturation into contractile cardiomyocytes (**Fig. 4A–B**). Constitutive phosphatase or viral-decoy expression during differentiation might otherwise disrupt lineage fidelity or stress-response signaling. The inducible design therefore provides both temporal control and physiological precision, ensuring translational activation occurs only when desired.

The improvements observed here complement extensive biochemical optimization of mammalian cell-free protein synthesis (CFPS) systems. Previous studies (11, 12, 28, 62, 63) demonstrated that extract activity depends on ionic composition, energy-regeneration systems, and the stability of initiation factors. Our genetic approach acts upstream of these parameters by stabilizing the translation machinery before lysis. Whereas biochemical tuning enhances reaction chemistry, pre-emptive removal of inhibitory phosphorylation ensures that extracts enter translation in a permissive, active state—together defining a dual genetic-biochemical strategy for human CFPS optimization.

By eliminating or minimizing eIF2α phosphorylation, the protocols presented here will allow exploration of the effects of the mRNA template and other cell type-specific factors. The EMCV or HCV IRESes based reporters used in this study provide a robust benchmark but requires specific IRES trans-acting factors (ITAFs) (e.g., PTB or PCBP1/2) (64, 65), whose abundance varies between cell types and differentiation states. Such variation may contribute to the extract-dependent translation efficiency we observed. Incorporating alternative 5′ leaders—cap-dependent 5’-UTRs, CrPV IRESs, or synthetic sequences designed for human translation—may yield new insights into translation regulation in different lysates. In addition, cell-type–specific differences in translation-associated factors, including initiation factors and RNA-binding proteins such as poly(A)-binding protein (PABPC1), may also contribute to variable translation efficiency. PABPC1 concentration and activity are known to be dynamically regulated by poly(A)-tail length, particularly in cardiomyocytes, where this mechanism fine-tunes translational output during growth and stress adaptation (66). For example, whereas IRES-dependent translation in cardiomyocyte extracts was robust, we were unable to detect translation of the m^7^G-capped *HBB* 5’-UTR Nanoluciferase reporter. Such variability in PABPC1 or related factors should be considered when comparing lysates from distinct human cell types or planning future optimization of IVT reactions. Translation efficiency is also influenced by tRNA availability and modification. Cell extracts from Expi293F, WI-38, and iPSC-derived cardiomyocytes likely differ in their pools of functionally active or modified tRNAs. Codon usage, tRNA charging levels, and wobble-base modification therefore represent additional axes for mRNA optimization. Integrating quantitative tRNA-omics with cell-type-specific lysate profiling could help identify limiting tRNAs and guide rational codon re-engineering to further enhance yields. Our findings establish a molecular foundation upon which these kinds of biochemical optimizations and interrogations can be layered, bridging genetic engineering with extract chemistry and paving the way toward robust, scalable, and physiologically relevant human translation platforms.

## Acknowledgments

We are grateful to Dr. Hanqin Li for iPSC cultivation guidance, and to the Doudna, Urnov, Hockemeyer, Ingolia and Braverman laboratories for generously sharing plasmids. We are grateful to Dr. Yumi Koga and Dr. Conner J. Langeberg for their critical reading of the manuscript and valuable suggestions.

## Author contributions

*Nikolay A. Aleksashin* - Conceptualization, Data curation, Formal analysis, Investigation, Methodology, Project administration, Supervision, Validation, Visualization, Writing – original draft, Writing – review and editing; *Rohan R. Shelke* - Formal analysis, Investigation, Methodology, Writing – review and editing; *Tianhao Yin* - Formal analysis, Investigation, Methodology, Writing – review and editing; *Jamie H. D. Cate* - Conceptualization, Formal analysis, Funding acquisition, Investigation, Methodology, Project administration, Supervision, Writing – original draft, Writing – review and editing.

---

**Figure S1.**
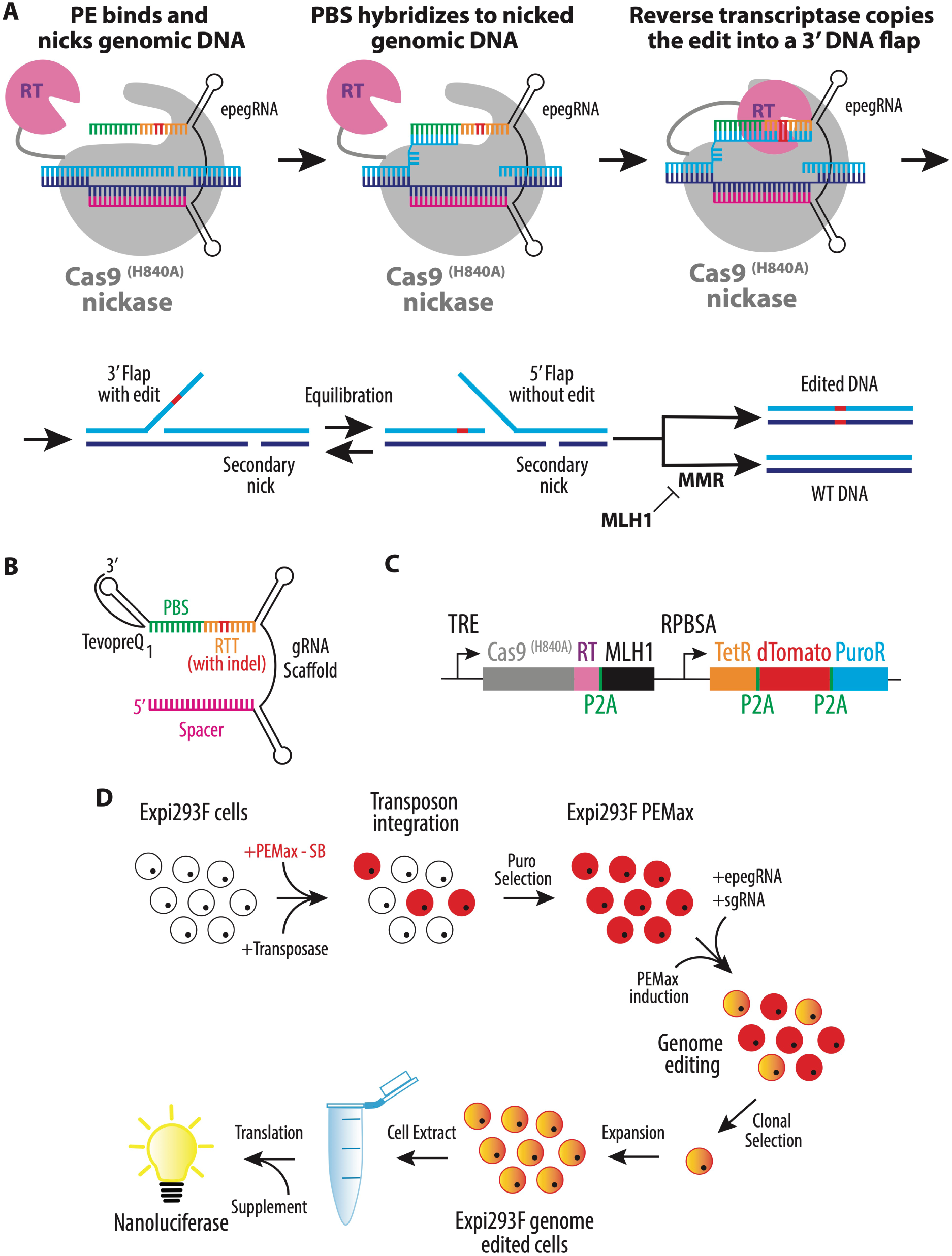
Prime-editing strategy for generating phospho-null eIF2α-S52A and eEF2-T57A Expi293F cell lines. **(A)** Mechanism of prime editing (PE). The Cas9(H840A) nickase fused to a reverse transcriptase (RT) introduces the desired sequence change without generating a double-strand break. The PE complex nicks genomic DNA guided by an engineered prime-editing guide RNA (epegRNA). The primer-binding site (PBS) of the epegRNA anneals to the nicked strand, allowing reverse transcription of the encoded edit into a 3′ DNA flap. Flap equilibration and mismatch repair (MMR) pathways resolve the edited and unedited strands to produce permanent sequence incorporation. To bias repair toward the edited strand and suppress degradation of the edited flap, the system includes a dominant-negative MLH1 (MLH1dn) domain that transiently inhibits MMR activity, thereby increasing prime-editing efficiency and product purity. A secondary nick on the non-edited strand further promotes repair in favor of the newly synthesized, edited DNA. **(B)** Structure of the engineered pegRNA (epegRNA) used in this study. The epegRNA includes a reverse-transcription template (RTT) encoding the desired nucleotide change, a PBS for priming, and a structured TevopreQ1 RNA aptamer extension at the 3′ end that protects against exonuclease degradation and improves editing efficiency. **(C)** Design of the inducible prime-editing vector (PEmax–SB). The Sleeping Beauty transposon encodes the Cas9(H840A)–RT fusion (PEmax) under a TRE promoter, followed by a self-cleaving P2A–mCherry–P2A–Puromycin resistance (PuroR) cassette. The RT module incorporates the MLH1dn domain to inhibit mismatch repair and stabilize the edited strand. The entire cassette is flanked by inverted repeats (IRs) for Sleeping Beauty transposition. **(D)** Workflow for generation of genome-edited Expi293F suspension cells. Expi293F cells were co-nucleofected with PEmax–SB and Sleeping Beauty transposase-encoding plasmids, followed by puromycin selection to obtain stable PEmax-expressing cells. These cells were nucleofected with plasmids encoding epegRNA and the corresponding nicking sgRNA to introduce the desired eIF2α S52A or eEF2 T57A edits. After doxycycline induction of PEmax, editing occurred in a controlled, transient window to minimize stress. Edited clones were expanded, validated by sequencing, and used for NanoLuc-based *in vitro* translation (IVT) assays to quantify translational yield.

**Figure S2.**
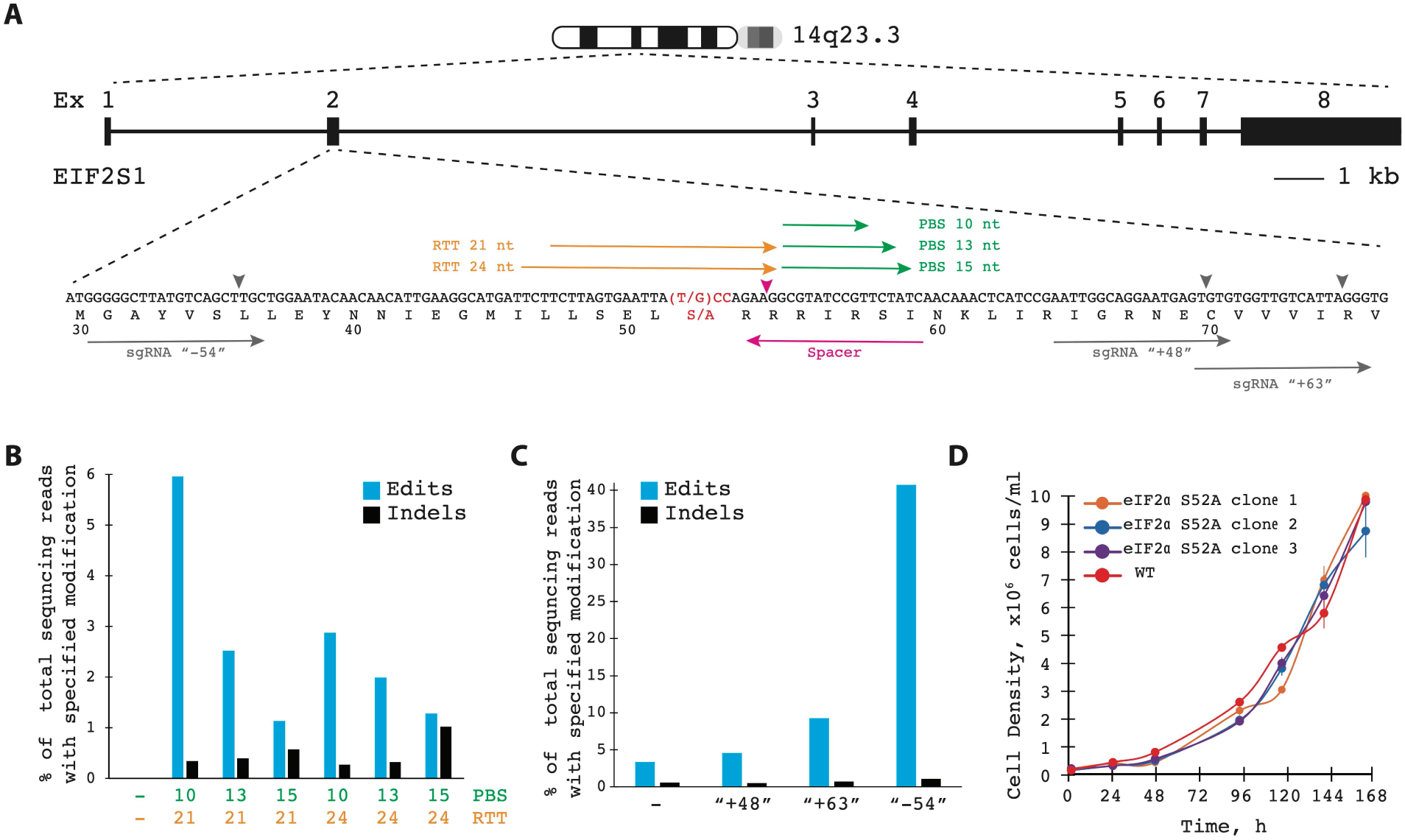
Optimization and validation of prime-editing strategy for introducing the eIF2α S52A mutation in Expi293F cells. **(A)** Schematic of the *EIF2S1* (eIF2α) genomic locus and prime-editing design. Exon 2 of *EIF2S1* was targeted to substitute Ser52 → Ala (TCC → GCC; red). The diagram shows the exon–intron structure, chromosomal location (14q23.3), and the positions of tested pegRNAs containing different primer-binding site (PBS) and reverse-transcription template (RTT) lengths (10–15 nt PBS; 21–24 nt RTT). The orientations of candidate nicking sgRNAs (“–54”, “+48”, “+63”) are indicated. Sequence is according to Homo sapience reference genome assembly GRCh38.p14 (GenBank assembly accession: GCA_000001405.29). **(B)** Editing efficiency as a function of PBS and RTT lengths. Bar graph shows the percentage of sequencing reads containing the precise edit (blue) or indels (black) across combinations of PBS and RTT designs. The 13-nt PBS / 24-nt RTT combination produced the highest editing frequency with minimal indels (< 3 %). **(C)** Comparison of nicking-sgRNA positions. Among tested secondary-nick locations, the “–54” sgRNA achieved the most efficient and accurate incorporation of the S52A substitution, confirming its suitability for subsequent clone generation. **(D)** Growth characteristics of genome-edited Expi293F eIF2α-S52A clones. Three independent S52A clones (orange, blue, purple) exhibit proliferation rates comparable to wild-type (red) Expi293F cells, confirming that the Ser52 → Ala substitution does not affect cellular fitness or suspension-culture viability.

**Figure S3.**
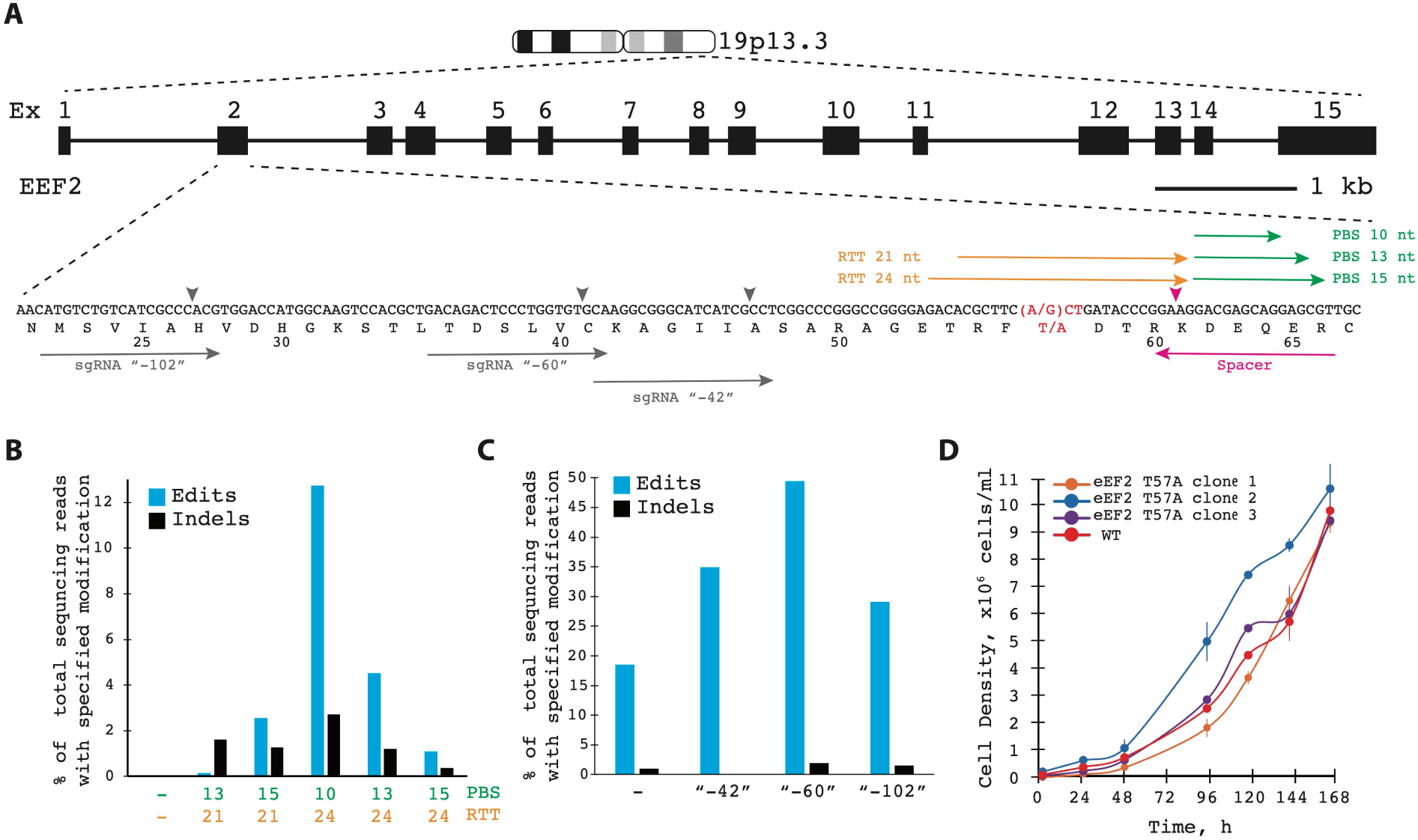
Optimization and validation of the eEF2 T57A prime-editing strategy in Expi293F cells. **(A)** Schematic of the *EEF2* genomic locus and prime-editing design. Exon structure of *EEF2* is shown with chromosomal location (19p13.3). Prime editing was used to introduce a Thr57 → Ala substitution (A → G, red) within exon 2. The orientations of tested nicking sgRNAs (“–42”, “– 60”, “–102”) and the lengths of reverse-transcription templates (RTT, 21–24 nt) and primer-binding sites (PBS, 10–15 nt) are indicated. Sequence is according to Homo sapiens reference genome assembly GRCh38.p14 (GenBank assembly accession: GCA_000001405.29). **(B)** Optimization of PBS and RTT lengths. Editing efficiency was quantified as the fraction of sequencing reads containing the precise substitution (blue) or indels (black). A 10-nt PBS / 24-nt RTT combination provided the highest editing yield with minimal indel formation. **(C)** Comparison of the efficiency of different nicking-sgRNA. Among the three tested secondary-nick sites, the “–60” yielded the highest frequencies of accurate editing, while the “–102” site showed lower efficiency. **(D)** Growth analysis of Expi293F eEF2-T57A clones. Three independent T57A clones exhibit proliferation kinetics indistinguishable from wild-type (WT) Expi293F cells, confirming that the mutation does not impair cell viability or suspension-culture growth.

**Figure S4.**
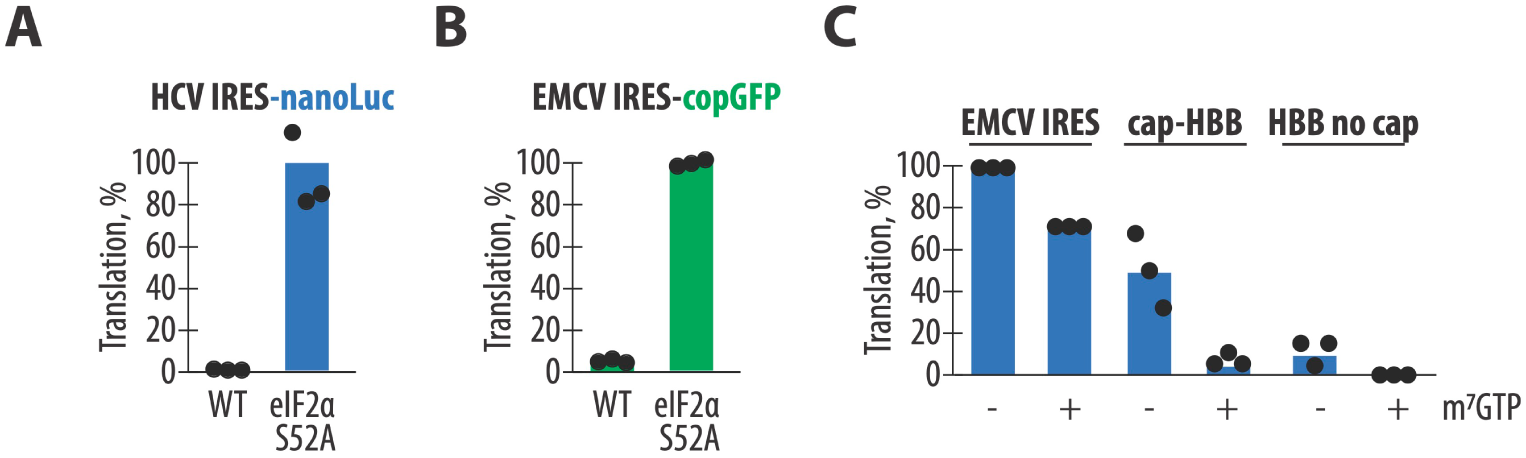
Translational activation in eIF2α S52A Expi293F cell extract is independent of reporter identity or initiation mechanism. (**A**) Translation of *NanoLuc* driven by the HCV IRES and (**B**) translation of copGFP driven by the EMCV IRES in wild-type and eIF2α S52A Expi293F cell extract. (**C**) Translation of EMCV-IRES NanoLuc mRNA, capped *HBB* (human β-globin) 5’UTR NanoLuc mRNA, and uncapped *HBB* reporter mRNA in eIF2α S52A cell extract in the absence or presence of cap analog m⁷GTP. All experiments were performed in biological triplicate.

**Figure S5.**
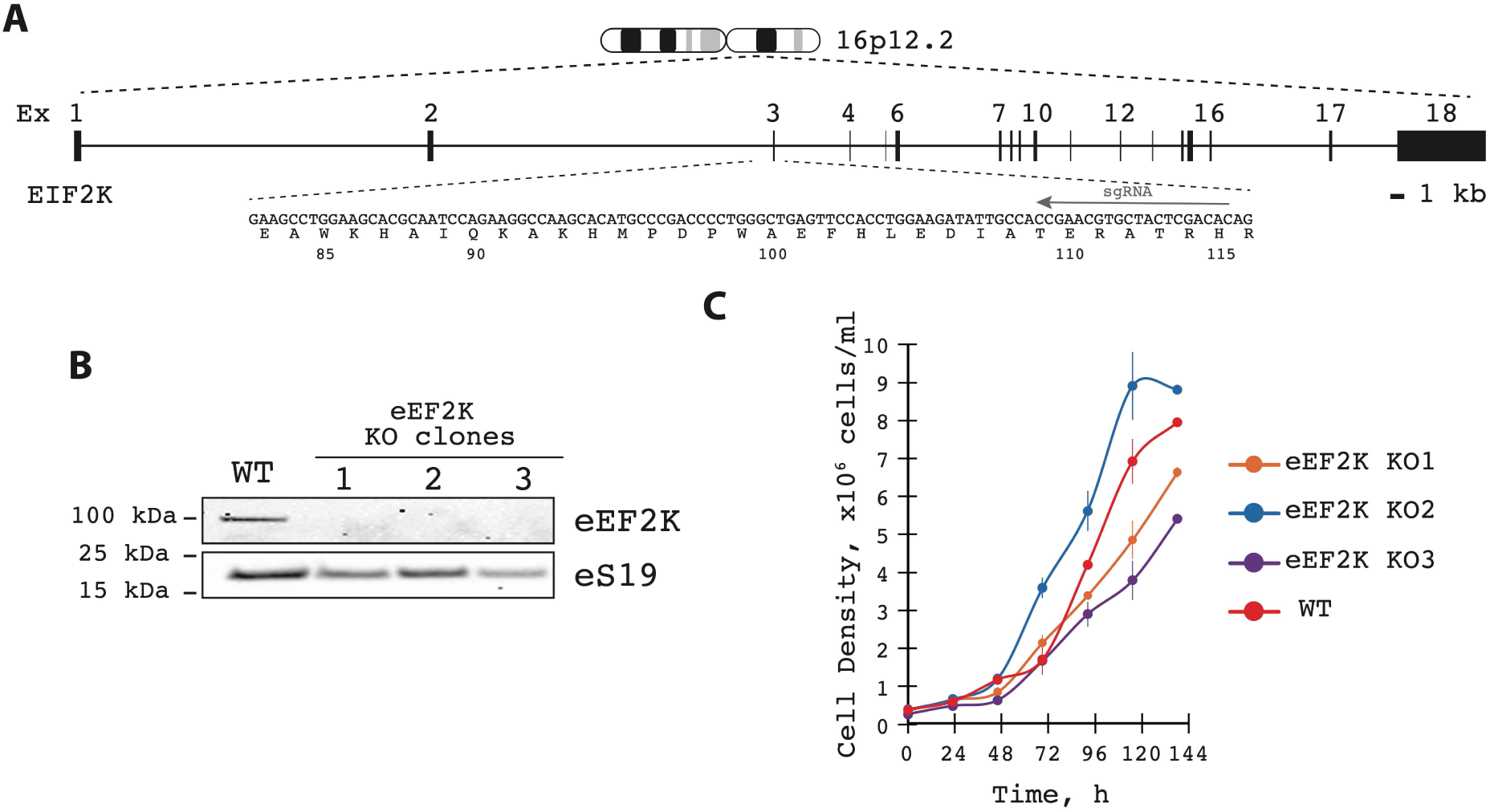
Generation and validation of eEF2K knockout Expi293F cell lines. **(A)** Schematic of the *EIF2K* (encoding eEF2 kinase) genomic locus and CRISPR–Cas9 targeting strategy. Exons and intron structure are shown with the chromosomal position (16p12.2). A single guide RNA (sgRNA) was designed to target exon 3, introducing a frameshift mutation predicted to disrupt kinase catalytic function. Sequence is according to Homo sapience reference genome assembly GRCh38.p14 (GenBank assembly accession: GCA_000001405.29). **(B)** Validation of eEF2K knockout clones by immunoblotting. Whole-cell lysates from three independent eEF2K-KO clones and wild-type (WT) Expi293F cells were analyzed by Western blot using an anti-eEF2K antibody. All KO clones showed complete loss of eEF2K protein, while eS19 served as a loading control. Gels are representative of two independent experiments. **(C)** Growth characteristics of eEF2K-KO clones. All three knockout clones proliferated at rates comparable to WT Expi293F cells, indicating that eEF2K loss does not affect suspension-culture viability or growth kinetics.

**Figure S6.**
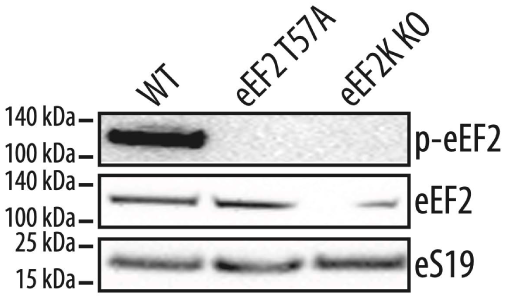
Mutation of eEF2 Thr57 and knockout of eEF2K eliminate eEF2 phosphorylation. Western blot analysis of extracts prepared from WT, eEF2 T57A mutant, and eEF2K knockout Expi293F cells. Immunoblotting with phospho-specific antibodies shows that eEF2 phosphorylation is completely abolished in both mutant and knockout strains, confirming loss of eEF2K-dependent modification at Thr57. Total eEF2 levels remain unchanged across all samples, as detected by pan-eEF2 antibody. Ribosomal protein eS19 serves as a loading control. Gels are representative of two independent experiments.

**Figure S7.**
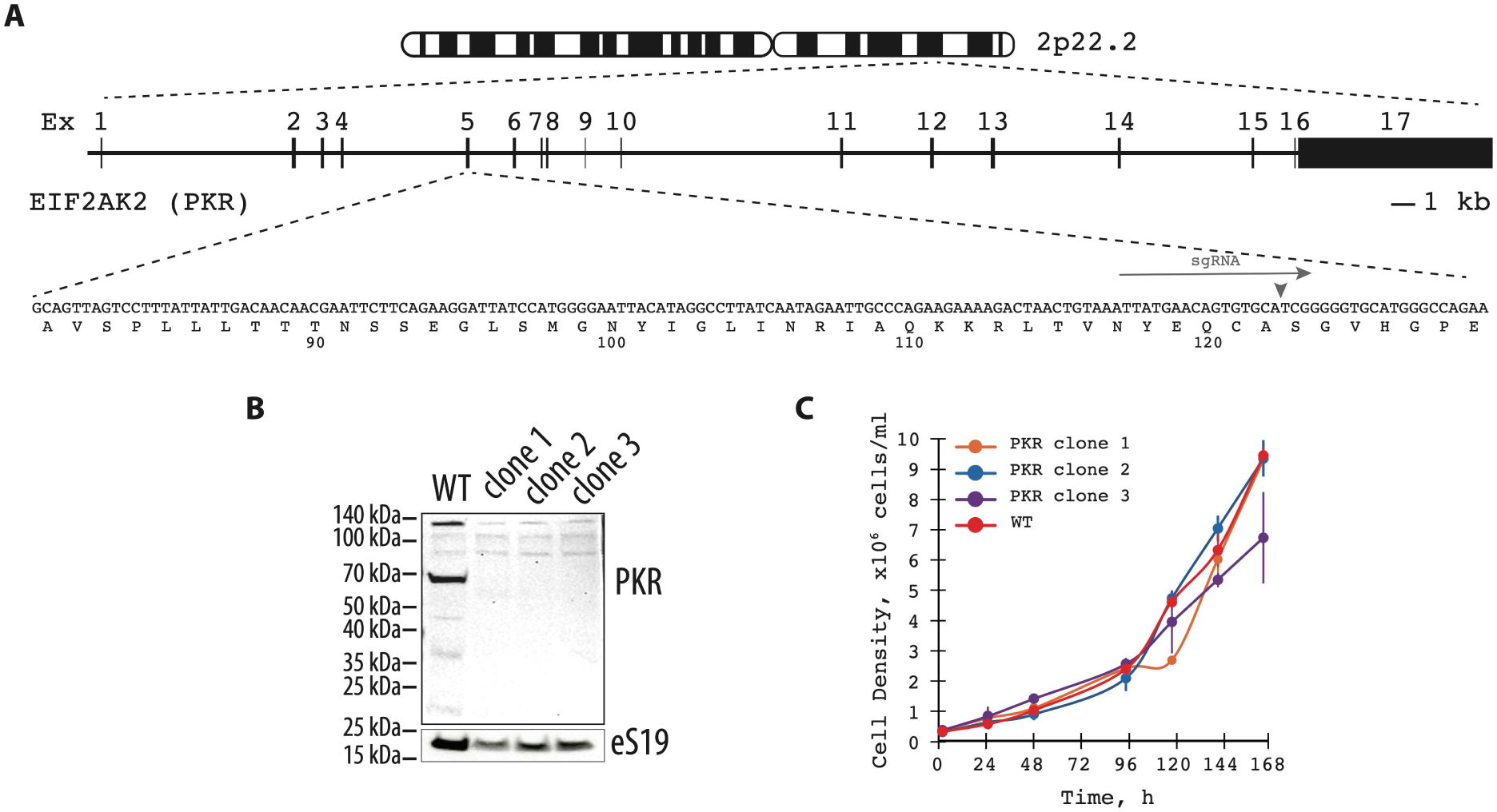
Generation and validation of PKR knockout Expi293F cells. (**A**) Schematic representation of the EIF2AK2 (PKR) gene locus on chromosome 2p22.2 showing exon organization and the sgRNA target site used for CRISPR/Cas9-mediated knockout. The sequence of the targeted region and corresponding amino acid positions are indicated below. Sequence is according to Homo sapience reference genome assembly GRCh38.p14 (GenBank assembly accession: GCA_000001405.29). (**B**) Western blot analysis of wild-type (WT) and three independent PKR knockout Expi293F clones. Immunoblotting with anti-PKR antibody confirms complete loss of PKR expression in all knockout clones, while ribosomal protein eS19 serves as a loading control. Gels are representative of two independent experiments. (**C**) Growth curves of WT and PKR knockout Expi293F clones cultured under standard suspension conditions. The proliferation rates of PKR knockout lines are comparable to WT, indicating that PKR is dispensable for cell viability and growth under non-stress conditions.

**Figure S8.**
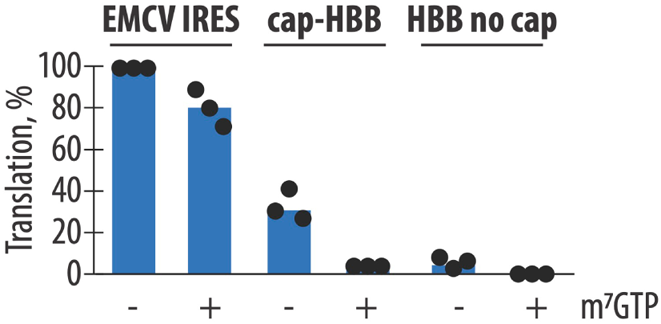
Both IRES-driven and cap-dependent nLuc reporters translate efficiently in fibroblast IVT extracts supplemented with GADD34/K3L. Normalized nanoLuciferase activities obtained using EMCV IRES–nLuc, capped *HBB*–nLuc, and uncapped *HBB*–nLuc reporters in the human fibroblast–based *in vitro* translation system after GADD34Δ /K3L overexpression. Reactions were carried out in the absence (–) or presence (+) of m⁷GTP cap competitor. Each dot represents an independent IVT reaction; bars show the mean.

**Figure S9.**
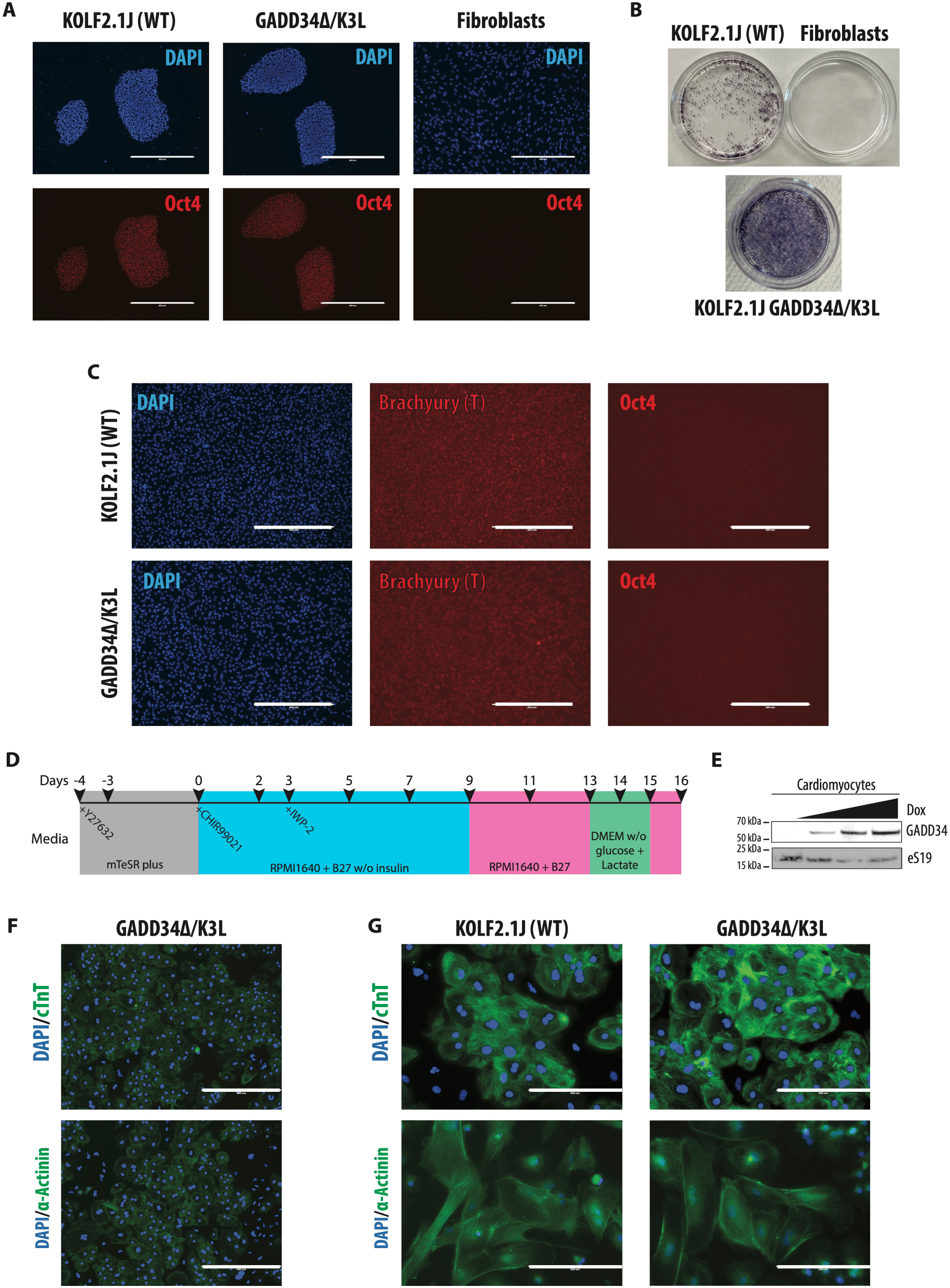
Characterization of iPSC pluripotency and cardiomyocyte differentiation. (**A**) Immunofluorescence analysis of pluripotency marker OCT4 (red) in wild-type KOLF2.1J iPSCs, GADD34Δ/K3L iPSCs, and human fibroblasts. DAPI (blue) marks nuclei. Scale bars are 400 µm. (**B**) Cytochemical alkaline phosphatase staining of undifferentiated KOLF2.1J and GADD34Δ/K3L iPSCs, confirming retention of pluripotent marker and loss of staining in differentiated fibroblasts. (**C**) Directed mesoderm differentiation assay used to validate trilineage potential. After 48 h of induction cells expressed the mesoderm marker Brachyury (T) and lost OCT4 expression, indicating successful commitment to the mesoderm lineage. Scale bars are 400 µm. (**D**) Schematic timeline of cardiomyocyte differentiation from human KOLF2.1J iPSCs using small-molecule Wnt-signaling modulation. Cells were maintained in mTeSR Plus with Y-27632 for 3–4 days before induction. Differentiation was initiated on day 0 with CHIR99021 in RPMI1640 + B27 without insulin, followed by IWP-2 treatment on day 3. On day 9, cultures were switched to RPMI1640 + B27 with insulin for maturation; metabolic selection (DMEM without glucose + 4 mM lactate) was performed on days 13–15. (**E**) Immunoblot analysis of doxycycline-inducible GADD34Δ expression in differentiated cardiomyocytes. Increasing doxycycline concentrations (0–1 µg/mL) induced dose-dependent accumulation of GADD34Δ protein. Ribosomal protein eS19 serves as loading control. Representative of two independent experiments. (**F–G**) Immunofluorescence staining of iPSC-derived cardiomyocytes for cardiac markers cTnT and α-actinin (green) with DAPI nuclear counterstain (blue). (**F**) GADD34Δ/K3L cells imaged at low magnification showing high differentiation efficiency. Scale bars are 400 µm. (see also corresponding wild-type control in Figure 4B). (**G**) Comparison of wild-type and GADD34Δ/K3L cardiomyocytes sarcomeric organization at higher magnification (scale bar 200 µm).

