## Supplementary material for "Overcoming the eIF2α Brake in Human Cell-Derived Translation Systems": Table S1

**Table S1. Oligonucleotides used for epegRNA and sgRNA plasmid cloning**

| Category | Name / Target | Sequence (5′–3′) | Purpose / Use |
|---|---|---|---|
| **PCR amplification primers** | epegRNA vector open F | GGTGTTTCGTCCTTTCCACAAGATA | PCR amplification of epegRNA and sgRNA backbones |
| | epegRNA vector open R | CGCGGTTCTATCTAGTTACGCGTTA | PCR amplification of epegRNA vector |
| | sgRNA vector open R | GTTTTAGACCTAGAAATAGCAAGTT | PCR amplification of sgRNA vector |
| **eIF2α epegRNA variable oligos** | eIF2α 10/21 | GCACCGAGTCGGTGCTCTTAGTGAATTAGCCAGAAGGCGTATCCGTCGCGGTTCTATCTAGTTAC | epegRNA variant cloning |
| | eIF2α 13/21 | GCACCGAGTCGGTGCTCTTAGTGAATTAGCCAGAAGGCGTATCCGTTCTCGCGGTTCTATCTAGTTAC | epegRNA variant cloning |
| | eIF2α 15/21 | GCACCGAGTCGGTGCTCTTAGTGAATTAGCCAGAAGGCGTATCCGTTCTATCGCGGTTCTATCTAGTTAC | epegRNA variant cloning |
| | eIF2α 10/24 | GCACCGAGTCGGTGCTCTTCTTAGTGAATTAGCCAGAAGGCGTATCCGTCGCGGTTCTATCTAGTTAC | epegRNA variant cloning |
| | eIF2α 13/24 | GCACCGAGTCGGTGCTCTTCTTAGTGAATTAGCCAGAAGGCGTATCCGTTCTCGCGGTTCTATCTAGTTAC | epegRNA variant cloning |
| | eIF2α 15/24 | GCACCGAGTCGGTGCTCTTCTTAGTGAATTAGCCAGAAGGCGTATCCGTTCTATCGCGGTTCTATCTAGTTAC | epegRNA variant cloning |
| **eEF2 epegRNA variable oligos** | eEF2 13/21 | GCACCGAGTCGGTGCACGCTTCGCTGATACCCGGAAGGACGAGCAGGAGCGCGGTTCTATCTAGTTAC | epegRNA variant cloning |
| | eEF2 15/21 | GCACCGAGTCGGTGCACGCTTCGCTGATACCCGGAAGGACGAGCAGGAGCGCGCGGTTCTATCTAGTTAC | epegRNA variant cloning |
| | eEF2 10/24 | GCACCGAGTCGGTGCGACACGCTTCGCTGATACCCGGAAGGACGAGCAGCGCGGTTCTATCTAGTTAC | epegRNA variant cloning |
| | eEF2 13/24 | GCACCGAGTCGGTGCGACACGCTTCGCTGATACCCGGAAGGACGAGCAGGAGCGCGGTTCTATCTAGTTAC | epegRNA variant cloning |
| | eEF2 15/24 | GCACCGAGTCGGTGCGACACGCTTCGCTGATACCGGGAAGGACGAGCAGGAGCGCGCGGTTCTATCTAGTTAC | epegRNA variant cloning |

| Category | Name / Target | Sequence (5′–3′) | Purpose / Use |
|---|---|---|---|
| **Constant oligos** | eIF2α epeg protospacer | TGGAAAGGACGAAACACCGATAGAACGGATACGCCTTCGTTTTAGACGTAGAAATAGCAA | Protospacer for eIF2α epegRNA constructs |
| | eEF2 epeg protospacer | TGGAAAGGACGAAACACCACGCTCCTGCTCGTCCTTCCGTTTTAGAGCTAGAAATAGCAA | Protospacer for eEF2 epegRNA constructs |
| | eIF2α/eEF2 epeg scaffold oligo | GCACCGACTCGGTGCCACTTTTTCAAGTTGATAACGGACTAGCCTTATTTTAACTTGCTATTTCTAGCTCTAAAAC | epegRNA scaffold oligo |
| **eIF2α sgRNA variable oligos** | eEF2 sgRNA −102 | TGGAAAGGACGAAACACCCATGTCTGTCATCGCCCACGGTTTTAGAGCTAGAAATAGCAA | sgRNA variant cloning |
| | eEF2 sgRNA −60 | TGGAAAGGACGAAACACCGACAGACTCCCTGGTGTGCAGTTTTAGAGCTAGAAATAGCAA | sgRNA variant cloning |
| | eIF2 sgRNA −54 | TGGAAAGGACGAAACACCGGGGGCTTATGTCAGCTTGCGTTTTAGAGCTAGAAATAGCAA | sgRNA variant cloning |
| | eEF2 sgRNA −42 | TGGAAAGGACGAAACACCCAAGGCGGGCATCATCGCCTGTTTTAGAGCTAGAAATAGCAA | sgRNA variant cloning |
| | eIF2 sgRNA +48 | TGGAAAGGACGAAACACCAATTGGCAGGAATGAGTGTGGTTTTAGAGCTAGAAATAGCAA | sgRNA variant cloning |
| | eIF2 sgRNA +63 | TGGAAAGGACGAAACACCGTGTGTGGTTGTCATTAGGGGTTTTAGAGCTAGAAATAGCAA | sgRNA variant cloning |
